# Modeling disorder, secondary structure formation, and amyloid growth in FG-nucleoporins

**DOI:** 10.64898/2026.04.07.716937

**Authors:** Maurice Dekker, Savio Ming Hou Chen, Vasista Adupa, Patrick R. Onck

## Abstract

Nuclear transport relies on intrinsically disordered FG-nucleoporins (FG-Nups) that form a dynamic selective barrier, yet experiments show that these proteins can also adopt highly ordered fibrillar structures. Capturing this duality within a single computational framework remains a major challenge. Here, we introduce 2BPA-HB, a sequence-resolved coarse-grained model that combines explicit, directional backbone hydrogen bonding with residue-specific side chain interaction centers derived from all-atom simulations. This design enables microsecond-scale simulations of the transitions between disordered, condensed, and amyloid-like states while maintaining computational efficiency. We first validate the model by simulating several experimentally resolved Nup98 fibril polymorphs, which retain their characteristic β-sheet registry and protofibril architecture. Seeded growth simulations further show that disordered FG fragments align with and extend these fibrillar templates in a sequence-specific manner, and can initiate secondary nucleation events on fibril surfaces. Applying the model to yeast FG-Nups, we find that cohesive FG domains form stable condensates with interaction networks dominated by FG motif contacts, consistent with experimental observations and prior simulations. The increased structural resolution of 2BPA-HB additionally reveals transient α- and β-structures within these condensates. Together, these results demonstrate that 2BPA-HB captures both liquid-like and fibrillar behavior of intrinsically disordered proteins within a single coarse-grained framework. The model provides a platform to study how disorder, secondary structure formation, and aggregation coexist in nuclear pore biology, and represents a step forward to a transferable coarse-grained approach for low-complexity proteins that undergo both phase separation and amyloid formation, as implicated in neurodegenerative diseases.

## Introduction

Protein aggregation is a biological process in which soluble proteins misfold and assemble into insoluble, often toxic, aggregates [1, 2]. This phenomenon is associated with various neurodegenerative disorders, including Alzheimer’s, Parkinson’s, and Huntington’s diseases. In these conditions, proteins such as amyloid-beta, alpha-synuclein, tau, and huntingtin undergo abnormal conformational changes that promote the formation of β-sheet-rich structures, which can stack into amyloid fibrils or other aggregate structures [1, 3]. These aggregates disrupt cellular functions and are implicated in the progressive neuronal damage characteristic of these diseases [1, 3]. An important feature of the aggregation process is secondary nucleation, where existing aggregates catalyze the formation of new aggregates, creating a self-reinforcing cascade [4–6]. Maintaining a disordered, soluble state is therefore critical in many biological systems, as transitions to ordered aggregates are often directly linked to pathology.

One biological process that heavily relies on a disordered, multivalent, and dynamic state is nuclear transport, where intrinsically disordered proteins form a selective permeability barrier that regulates molecular traffic between the nucleus and cytoplasm [7]. This barrier is maintained by intrinsically disordered proteins known as FG nucleoporins (FG-Nups), which contain regions of hydrophobic FG (phenylalanine–glycine) motifs interspersed with hydrophilic spacers, anchored to the inner surface of the nuclear pore complex (NPC) [8]. These FG motifs are crucial for the formation of a dynamic and selective barrier [9–11], enabling selective passage of macromolecules while preventing the uncontrolled diffusion of others [12].

Experimental studies have demonstrated that several FG-Nups can phase separate into liquid-like condensates that exhibit properties similar to the selective permeability barrier of the NPC [10, 13, 14]. In vitro studies show that these phase-separated condensates rapidly transition into hydrogels containing β-sheet-rich regions [15, 16]. Interestingly, FG-Nups have also been found to aggregate into amyloid fibrils under certain conditions [17–21]. These amyloids, characterized by highly ordered β-sheet arrangements, highlight the dual behavior of FG-Nups. While they support the permeability barrier in their phase-separated state, they also possess an intrinsic tendency to form amyloids, which could disrupt NPC function and impact cellular homeostasis [22]. The ability of FG-Nups to form amyloids outside the NPC raises important questions about the mechanisms that drive their aggregation and the cellular safeguards that normally prevent it [18, 23].

Molecular dynamics (MD) simulations have become an essential tool for studying the molecular mechanisms underlying phase separation and aggregation of intrinsically disordered proteins [24]. Atomistic MD has yielded valuable insights into sequence-dependent interactions, conformational ensembles, and the early stages of β-sheet formation in aggregation-prone systems [25–27]. However, atomistic simulations remain limited in their ability to access the large molecular assemblies and long timescales that characterize protein condensates, hydrogels, and amyloid fibrils. As a result, coarse-grained molecular dynamics (CGMD) models have become indispensable for bridging this gap [10, 28–32], enabling the study of mesoscale organization and phase behavior across a wide range of protein systems.

Most CGMD models developed for disordered proteins adopt a one-bead-per-residue representation. While these models are computationally efficient and successful in capturing global polymer properties and phase behavior [33], their reduced structural resolution limits their ability to describe the formation of secondary structure motifs. In particular, the emergence of β-sheets depends on directional backbone hydrogen bonding and anisotropic side chain interactions, which cannot be accurately represented when each amino acid is reduced to a single isotropic interaction site. Recent efforts to introduce more detailed coarse-grained descriptions [34–37], such as our 2BPA-Q model for polyglutamine aggregation [38], have demonstrated that adding explicit backbone and side chain degrees of freedom enables the formation of ordered β-structures in a computationally tractable manner. These advances highlight the need for higher-resolution coarse-grained models that retain the essential interactions required for secondary-structure formation while remaining amendable to large protein assemblies.

To investigate FG-Nup aggregation mechanisms in the context of the NPC, we present a CGMD model that extends our previous 2BPA-Q framework [38] from glutamine (Q) to all 20 amino acids. Although 2BPA-Q was originally designed to study polyglutamine aggregation, its two-beads-per-amino-acid (2BPA) representation is well suited to capture secondary structure formation in intrinsically disordered systems more broadly. By combining an increased level of coarse-grained resolution with explicit, directional backbone hydrogen bonding interactions, the model enables α-helices and β-sheets to emerge spontaneously. The inclusion of a separate side chain bead allows for sequence-specific packing and stabilization of ordered motifs, while the hydrogen bonding scheme remains unbiased, ensuring that secondary structure formation arises from the underlying interactions rather than imposed structural restraints.

Here, we generalize this approach to develop a sequence-resolved model tailored to FG-Nups, which we refer to as 2BPA-HB, reflecting its explicit treatment of backbone hydrogen bonding (HB) that enables β-sheet formation. Using this model, we successfully reproduce experimentally resolved fibril structures and characterize their stability and conformational dynamics. In addition, the 2BPA-HB framework enables the study of FG-Nup phase transitions, amyloid growth, and secondary nucleation, highlighting its ability to capture sequence-specific disorder-to-order processes. Together, these results demonstrate that 2BPA-HB provides a versatile and transferable coarse-grained approach for describing both disordered and amyloid-like states of FG-Nups, offering new insight into their aggregation behavior and implications for NPC function.

Beyond FG-Nups, the framework is applicable to other intrinsically disordered proteins that exhibit both phase separation and aggregation, offering new opportunities to study disorder-to-order transitions across a broad range of neurodegenerative diseases.

## Methods

### Design of the 2BPA-HB model

The 2BPA-HB model is an extension of the 2BPA-Q model, which we previously developed to study the aggregation behavior of polyglutamine [38]. Where the 2BPA-Q model focused specifically on glutamine, the 2BPA-HB model expands this approach to all 20 amino acids, making it applicable to a broad range of proteins. In this model, each amino acid is represented by two beads: one for the backbone (BB) and one for the side chain (SC); except for glycine, which is represented by a BB bead only.

The bonded potential for the backbone beads in the 2BPA-HB model is directly adopted from the 1BPA force field [39]. This potential is given by:

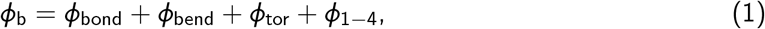

and includes terms for bond stretching, bending, torsion, and 1–4 coupling interactions. The bond stretching between two covalently bonded backbone beads is described by a stiff harmonic potential with an equilibrium distance of *b* = 0.38 nm and a force constant of 8038 kJ/nm^2^/mol. The bending and torsion potentials distinguish between glycine, proline, and all other residues, allowing the model to capture the distinct backbone stiffness characteristic of these amino acids, in contrast to many CGMD models that merely employ a uniform backbone stiffness. Additionally, the 1–4 distance potential (between beads *i* and *i* +3) ensures proper sampling of (*θ, α*)-space, addressing the uncoupled backbone dihedrals. For further details on the potentials in Eq. 1, we refer to Ghavami et al. [39], where these interactions were originally introduced as part of the 1BPA force field.

In the 2BPA-HB model, each side chain is assigned a residue-specific length and orientation relative to the protein backbone, described by parameters *a* and *b* (Fig. 1a). The orientation of the side chain bead is maintained through the introduction of a virtual particle at position 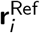, which serves as a reference point for the side chain’s position. This virtual particle has no nonbonded interactions and exists solely to stabilize the geometry of the side chain. For each residue, the position of the virtual reference point is defined as a fraction *a* along the vector between BB beads *i* − 1 and *i* +1 (Fig. 1a):

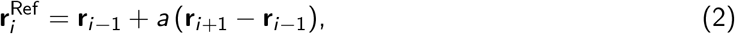

where *a* is a value between 0 and 1. The position of the side chain is restraint by a harmonic angle potential, with an equilibrium angle of 180^◦^ and force constant 1000 kJ/mol/rad^2^, between the side chain (SC_*i*_), backbone (BB_*i*_), and reference point (Ref_*i*_). This means that the orientation of the side chain is controlled by the parameter *a*, rather than by the harmonic angle itself, which is more stable for equilibrium angles close to 180 degrees. As a result of this construction, the side chain beads (SC_*i*_) remain in the plane defined by the backbone beads BB_*i*−1_, BB_*i*_, and BB_*i*+1_.

**Fig. 1.**
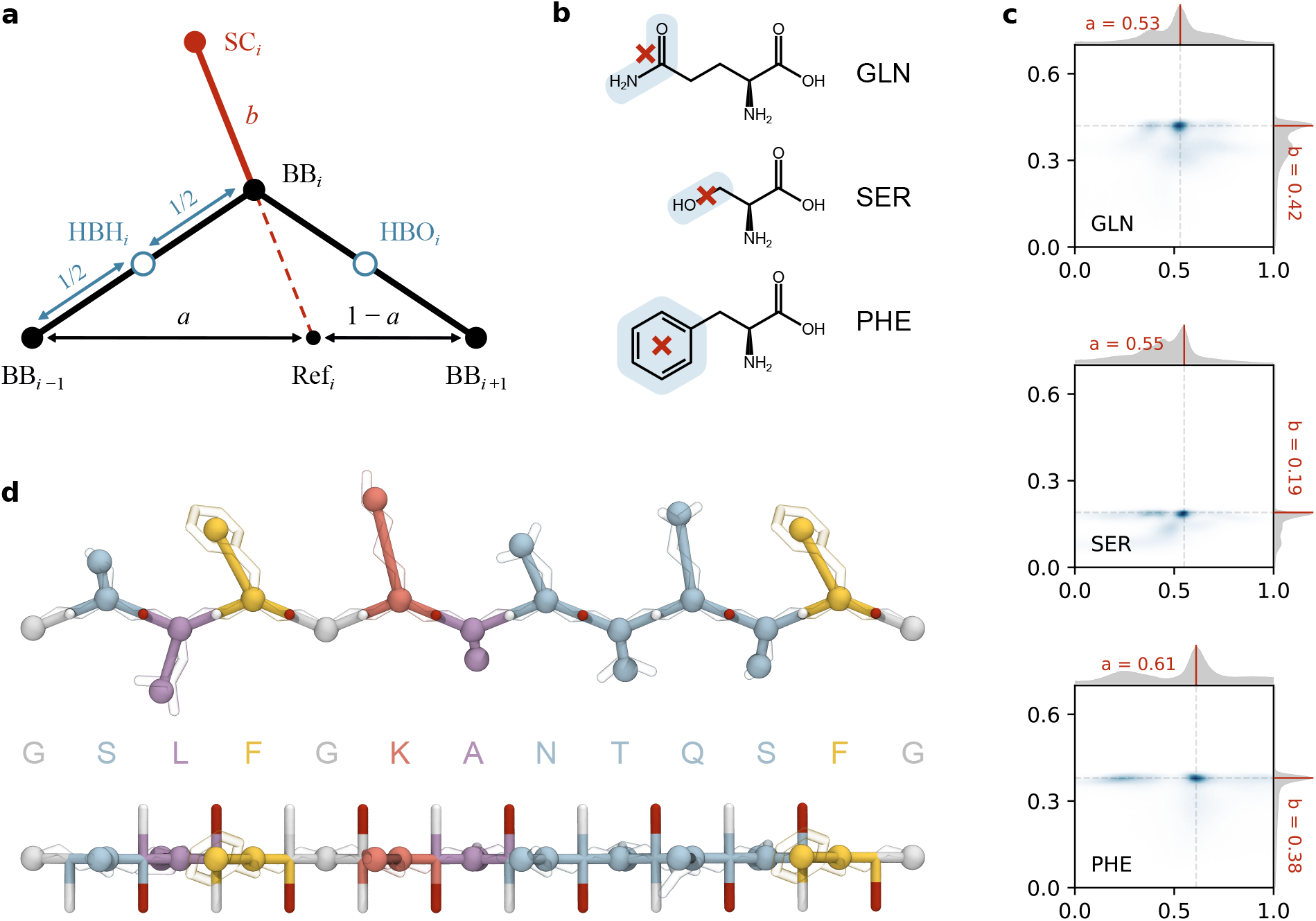
Design of the side chain geometry in the 2BPA-HB model. (**a**) Schematic illustration of the procedure used to position side chain (SC) beads. A reference point is placed at fractional distance *a* along the vector from BB_*i*−1_ to BB_*i*+1_, and the SC bead is positioned at distance *b* from BB_*i*_ along the reference–backbone direction. (**b**) Extraction of the geometric parameters *a* and *b* from all-atom simulations. For each residue type, the SC interaction site is defined as the center of geometry of the three heavy atoms farthest from the backbone (example: GLN; selected atoms highlighted in blue, interaction site marked with a red cross). For residues with fewer than three heavy atoms, all heavy atoms are used (example: SER), and for aromatic residues containing a benzene ring, the entire ring is selected (example: PHE). A complete list of selected atoms for each residue type is provided in Suppl. Table S1. (**c**) Two-dimensional distributions of extracted (*a; b*) values from all-atom trajectories. One-dimensional marginal distributions for *a* and *b* are shown above and to the right of the main plot. For each residue type, the most probable (*a; b*) pair is used as the side chain placement in the 2BPA-HB model. Full distributions for all amino acids are provided in Suppl. Fig. S1. (**d**) Example FG fragment represented in the 2BPA-HB model. The coarse-grained structure is overlaid with the corresponding all-atom configuration (heavy atoms only, shown in transparency) to illustrate the alignment between the atomistic side chain and the coarse-grained SC bead. Two views are shown, rotated by 90^◦^, highlighting that the hydrogen bonding beads (HBH, white; HBO, red) are oriented perpendicular to the plane defined by the BB and SC beads. Residues are colored by type: polar (blue), hydrophobic (purple), aromatic (yellow), cationic (red), and glycine (white).

This method, however, cannot place side chains on terminal residues at the end of the protein chains, as they lack the necessary neighboring backbone atoms (*i* − 1 or *i* + 1). To resolve this, an additional backbone bead is introduced at both termini, ensuring that the side chains at the ends of the sequence can be properly positioned.

### Derivation of side chain parameters

Side chain interactions play an important role in stabilizing amyloid fibrils [40, 41]. In these fibers, the close packing of side chains from neighboring protein strands leads to a “zipper-like” structure, where complementary side chains interlock along the fibril’s length. These interactions help maintain the highly ordered, β-sheet-rich architecture characteristic of amyloids, making side chain zippering a critical factor in the formation and stability of these aggregates.

To capture this side chain behavior in our coarse-grained model, we extracted side chain parameters from all-atom MD trajectories of multiple FG-Nup segments. A key step in this process is correlating the position of the atomistic side chain with the placement of the side chain (SC) bead in the 2BPA-HB model. While using the center-of-mass of the atomistic side chain might seem like a natural approach, this method did not yield stable amyloid fibrils with the correct geometry.

To reproduce polyQ amyloid fibrils with correct structural dimensions, the SC bead in the 2BPA-Q model was positioned farther from the backbone than, for example, the side chain’s center of mass. To generalize this approach to all amino acids, we placed the SC bead at the center of geometry of the three heavy atoms farthest from the backbone. For example, in glutamine, the SC bead is placed at the center of geometry of the three distal heavy atoms C_*δ*_, O_*ε*1_, and N_*ε*2_ (see Fig. 1b). For residues with fewer than three heavy atoms in the side chain, the center of geometry of the available atoms is used; for serine, this would be the atoms C_*β*_ and O_*γ*_ (see Fig. 1b). For residues with aromatic rings, all heavy atoms of the ring are included in the calculation. For example, the SC bead in phenylalanine is placed at the center of geometry of C_*γ*_, C_*δ*1_, C_*δ*2_, C_*ε*1_, C_*ε*2_, and C_*”*_ (see Fig. 1b). A complete list of interaction site definitions for each residue type is provided in Suppl. Table S1.

With the interaction centers defined, we extracted the SC parameters from all-atom trajectories of five FG-Nup segments: Nsp1 (AA 1–172), Nup145N (AA 1–242), Nup60 (AA 389–539), Nup98 (AA 85–124), and Nup98 (AA 165–204); see Supporting Information for details on the all-atom simulation procedure. Each trajectory spans at least 500 ns, with frames sampled every 0.1 ns. For each frame, the relevant SC parameters, denoted as *a* and *b*, were determined for every residue. This was done by projecting the SC bead position onto the plane formed by three consecutive backbone beads: the C_*a*_ atoms from the preceding residue, the current residue, and the following residue. From this projection, a pair of values (*a, b*) was extracted, capturing the relative positioning of the side chain.

All of these parameter pairs were combined into a large dataset for each amino acid type. Notably, some amino acids had multiple distinct conformations, resulting in a variety of (*a, b*) combinations. Given that the current implementation of the 2BPA model does not support dynamic side chains, we binned the data onto a 0.01 grid and selected the most frequently occurring (*a, b*) pair as the optimal SC parameter for each amino acid (Fig. 1c). Note that this procedure identifies the dominant joint conformation in (*a, b*) space, rather than independently selecting the most frequent values of *a* and *b*. An overview of the (*a, b*) distributions for all amino acids is provided in Suppl. Fig. S1.

### Nonbonded interactions

The 2BPA-HB model employs a hydrophobic potential consistent with the 1BPA model, represented by a shifted 8–6 Lennard-Jones potential:

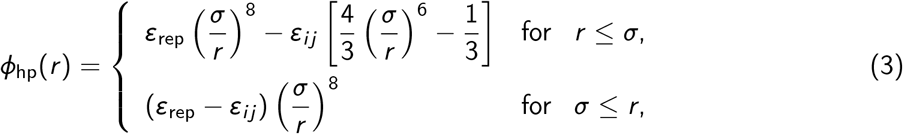

where *σ* = 0.476 nm is identical to the value used in the 2BPA-Q model [38]. This choice ensures that the combined excluded volume of the backbone and side chain beads approximately matches that of a single 1BPA bead [10, 42]. Moreover, this choice of *σ* yields a bead diameter that matches the characteristic interstrand spacing observed in β-sheets [41, 43]. Because this spacing is determined primarily by backbone geometry, which is identical for all amino acids, the same bead diameter can be used across residue types, allowing stable β-sheet formation without steric clashes. The interaction strength *ε*_*ij*_ for each pair of beads *i, j* is calculated using the combination rule:

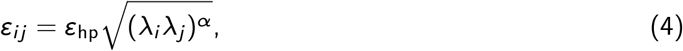

where λ_*i*_ is the relative hydrophobicity scale of bead *i*, with all 2BPA-HB λ_*i*_ values listed in Suppl. Table S1. The parameters *ε*_hp_ = 13 kJ/mol and *ε*_rep_ = 10 kJ/mol are constants taken from the 1BPA model [42], and the exponent *α* is set to 0.30.

As in our previous work [38], side chain interactions are represented by single coarse-grained beads interacting through isotropic attractive potentials. While this simplified description does not explicitly resolve the directionality of hydrogen bonds, it captures their overall cohesive contribution together with other solvation-driven and nonspecific interactions. Because experimental structures show that FG-Nup fibrils are stabilized in part by specific side chain hydrogen bonding interactions [21], we introduced a modest “polar boost” to approximately account for the stabilizing effect of these interactions at the coarse-grained level. Concretely, for polar residue pairs (N, Q, T, and S), the relative hydrophobicity scales λ_*i*_ are increased by 0.3 to enhance favorable polar–polar interactions:

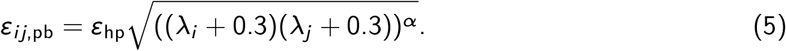

This adjustment enhanced secondary structure formation and improved the stability of the tertiary amyloid architecture throughout the simulations, while preserving the intrinsic disorder of individual monomers in solution.

To further enhance the stability of FG-Nup fibrils in our simulations, we incorporated a specific interaction potential between phenylalanine (F) side chains to mimic π–π interactions commonly observed between aromatic groups [20]. This interaction was inspired by the π–π potential used in the 1BPA-2.1 force field [31]. In our model, this phenylalanine-specific interaction follows a dedicated potential form to capture the energetic favorability of π–π stacking:

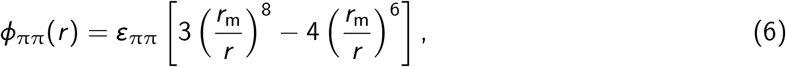

where *r*_m_ = 0.45 nm and *ε*_ππ_ = 5.0 kJ/mol. Additionally, to balance this added attractive energy, we adjusted the relative hydrophobicity scale λ_*i*_ for phenylalanine side chains, slightly lowering it to maintain the overall hydrophobic interaction profile of the residue.

In our model, electrostatic interactions between charged side chains are taken directly from the 1BPA force field and are modeled with a modified Coulomb potential:

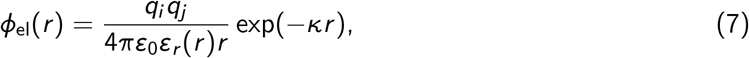

where *q*_*i*_ is the charge of residue *i*, and *ε*_0_ is the vacuum permittivity. We use a Debye screening coefficient of *k* = 1.27 nm^−1^ to simulate physiological salt conditions (150mM). Additionally, the dielectric screening effect of the solvent is described by a sigmoidal function:

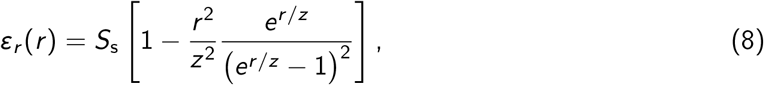

where *S*_s_ = 80 corresponds to the dielectric constant of water and *z* = 0.25 nm defines the range of the screening effect.

The hydrophobic interactions of charged side chains are modified to be neutral, effectively reducing repulsion between like-charged side chains. This modification, previously implemented in the 1BPA-2.1 force field [31], further improves fibril stability by balancing hydrophobic and electrostatic interactions without allowing excessive repulsion between similarly charged side chains.

### Hydogen bonding scheme

We implement a hydrogen bonding scheme that is similar to that used in the 2BPA-Q model, which is based on the approach of Chen and Imamura [44, 45]. In this scheme, backbone hydrogen bonds are modeled by adding two hydrogen bonding beads for each residue: one for the hydrogen donor and another for the oxygen acceptor. These beads are modeled as virtual interaction centers, positioned relative to three consecutive C_α_ backbone beads (Eqs. 9 and 10). The hydrogen bonding beads are placed at the midpoint of the backbone bond, at a distance of *ℓ*_HB_ = 0.238 nm perpendicular to the backbone bond (Fig. 1d). For proline residues, the atomistic side chain attaches directly to the nitrogen in the backbone, removing the hydrogen atom and thus the ability to act as a donor. Consequently, in the 2BPA-HB model, proline residues lack a virtual hydrogen bead, mirroring this structural constraint.

The placement of the HB beads differs from that in the 2BPA-Q model [38], where the HB beads were positioned at 1/3 and 2/3 of the backbone bond length, as originally proposed by Chen and Imamura [44]. Most amyloid fibrils, including FG-Nup fibrils [20], typically consist of parallel β-sheets. However, parallel β-sheets tend to be less stable than anti-parallel β-sheets, such as those found in polyQ amyloids [46], due to the suboptimal hydrogen bonding pattern in parallel arrangements. By adjusting the placement of the hydrogen bond bead to the midpoint of the backbone bond, we enhance the stability of amyloid fibrils that are characterized by parallel β-sheets.

The positions of the hydrogen bonding beads are constructed from the positions of the backbone beads, **r**_*i*_, using the following equations:

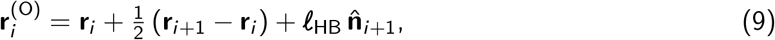

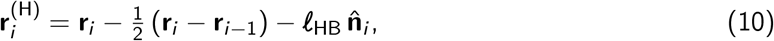

where

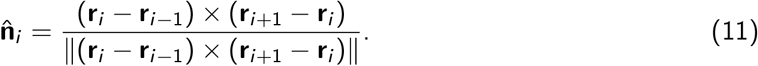

Here, *ℓ*_HB_ represents the perpendicular distance between the backbone bond and the hydrogen bonding bead, which is set at 0.238 nm to replicate the typical backbone-to-backbone separation of 4.76 Å found between adjacent β-strands [41, 43]. The hydrogen bonding interaction centers are implemented in GROMACS [47] using a customized 3OUT virtual site setup [38].

Interactions between hydrogen bonding beads are selectively applied to mimic hydrogen bonding. Hydrogen beads (donors) do not interact with each other, nor do oxygen beads (acceptors); only donor–acceptor interactions (O–H) are considered. The interaction between oxygen and hydrogen is modeled with a shifted Lennard-Jones potential:

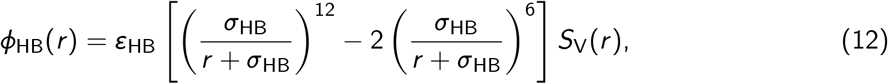

where *r* is the distance between the hydrogen and oxygen beads, *σ*_HB_ = 0.51 nm represents the interaction range, and *ε*_HB_ = 7.0 kJ/mol specifies the interaction strength. An ideal hydrogen bond configuration, characterized by *r* = 0, yields the maximal interaction energy of *ε*_HB_. To assign a clear physical meaning to the parameter *σ*_HB_ as the effective interaction diameter, the hydrogen bonding potential is multiplied by a potential-switch function [48],

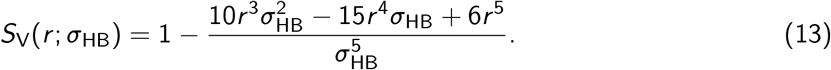

This construction ensures that the interaction smoothly decays to zero at *r* = *σ*_HB_, limiting the effective interaction range.

## Results

### Stability of FG-Nup fibrils in the 2BPA-HB model

To assess whether the 2BPA-HB model can reproduce the structural stability of experimentally observed FG-Nup assemblies, we first simulated pre-assembled FG-Nup fibrils using the 2BPA-HB model. Five experimentally determined structures were tested: four polymorphs of the Nup98_FG_(85–124) sequence (PDB IDs: 7Q64, 7Q65, 7Q66, 7Q67) [20] and one fibril of Nup98_FG_(298–327) (PDB ID: 8CI8) [21]. Each structure was simulated for 5 μs using fibril segments composed of 20, 30, or 50 layers to probe the length dependence of fibril stability. Here, a layer denotes one cross-β repeat along the fibril axis, consisting of one β-strand from each protofibril stacked in register. Thus, increasing the number of layers corresponds to extending the fibril length while preserving the lateral protofibril organization.

Across all systems, we monitored the root-mean-square deviation (RMSD) and total number of backbone hydrogen bonds over the simulation time as a measure of equilibration (Suppl. Figs. S2 and S3). The fibril length strongly influenced fibril stability. In systems composed of only 20 layers, not all polymorphs remained intact; instead, several exhibited progressive unraveling initiated by chain dissociation at the termini and followed by gradual disruption of the fibril core. In contrast, all fibrils with 30 or 50 layers reached a stable equilibrium state in which the core β-sheet architecture was preserved. Transient dissociation and re-association events were observed even in these larger systems, indicating a dynamic exchange between fibril-bound and free chains. This reversible behavior suggests that a minimal local concentration (or equivalently, a minimal fibril length) is required to maintain a stable fibril in the simulations.

As an illustrative example, we analyze polymorph 1 (PM1) of the Nup98_FG_(85–124) segment (Fig. 2). PM1 is the most abundant polymorph of the Nup98_FG_(85–124) fibril observed experimentally [20], and it also emerges as the most stable polymorph in our simulations. Corresponding analyses for the other polymorphs are provided in Suppl. Figs. S4–S7.

**Fig. 2.**
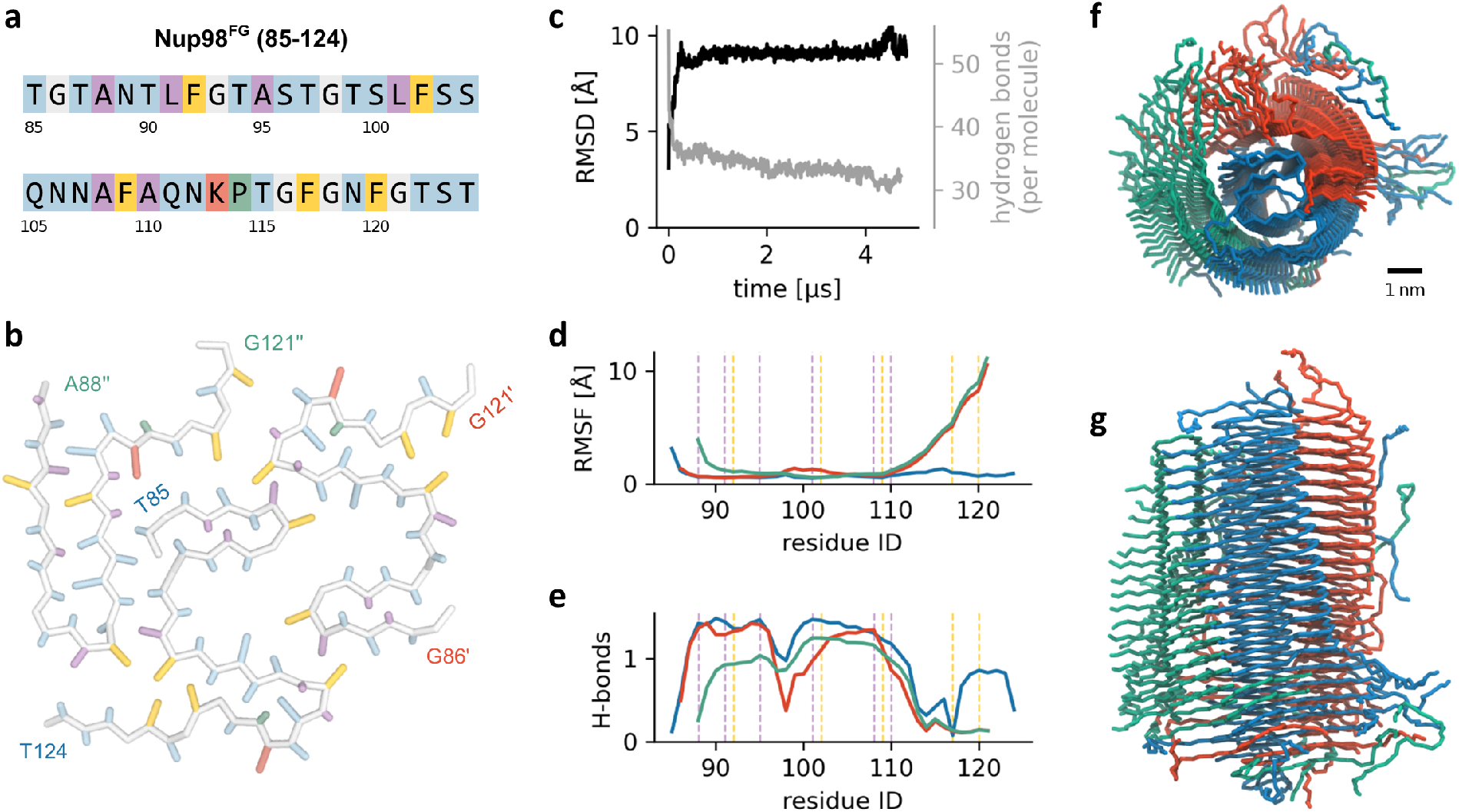
Stability and structural dynamics of the Nup98^FG^(85–124) PM1 fibril. **a** Amino acid sequence of Nup98^FG^(85– 124), colored by residue type: polar (blue), hydrophobic (purple), aromatic (yellow), cationic (red), glycine (white), and proline (green). **b** Single cross-section of the PM1 fibril (PDB ID: 7Q64), highlighting the side chain packing within the coarse-grained representation and the resulting organization of the three protofibrils. **c** Time evolution of the RMSD of the fibril core (black) and the average number of backbone hydrogen bonds per molecule (gray). Only the middle 10 layers were used for the RMSD calculation, as terminal chains transiently dissociate from the fibril ends. Because each non-proline residue can donate and accept one hydrogen bond, the total number of hydrogen bonds can exceed the number of residues. **d** Average RMSF of residues in each protofibril, calculated from the middle 10 layers and color coded as in panels (**f**) and (**g**). Note that the protofibrils do not have the exact same length as in the resolved structure some terminal residues were missing. Dashed lines indicate the positions of phenylalanine (yellow) and hydrophobic (purple) residues. **e** Average number of backbone hydrogen bonds per residue for each protofibril. The tertiary structure of the fibril core is stabilized by attractive side chain interactions, whereas the C-terminal regions of each protofibril show reduced hydrogen bonding. This reduced stability reflects both the absence of hydrophobic residues in these segments and electrostatic repulsion between the Lys113 residues. Only the blue protofibril has an increased stability at the C-terminal because of the enhanced number of hydrogen bonds. **f**,**g** Top (**f**) and side (**g**) views of the equilibrated 30-layer fibril at *t*\* = 5 µs, illustrating preservation of the overall protofibril architecture.

The PM1 fibril preserves its characteristic three–protofibril architecture throughout the simulation. Snapshots of the equilibrated structure (Fig. 2f,g) show that the global fold of the fibril is maintained, with only minor adjustments in the relative orientation of the protofibrils. To quantify structural stability, we computed the RMSD of the central 10 layers relative to the experimental structure (Fig. 2c). The RMSD increases during the initial relaxation phase and then plateaus at approximately ∼9 Å, indicating modest but stable conformational rearrangements while retaining the overall β-sheet organization of the fibril core. Most of this deviation originates from partial unraveling of the C-terminal segments (≈10 residues) in two protofibrils (red and green in Fig. 2d), whereas the blue protofibril retains its native fold along its entire length.

Consistent with the RMSD analysis, the total number of backbone hydrogen bonds reaches equilibrium after roughly 2 μs and stabilizes at approximately 33 hydrogen bonds per molecule (Fig. 2c). The fibril reaches equilibrium after ∼2 μs, stabilizing at around 33 hydrogen bonds per molecule. This average includes molecules that have partially or fully dissociated from the fibril, so the number of hydrogen bonds within the fibril core itself is higher. The plateau indicates that the fibril reaches a steady state in which the pattern of backbone hydrogen bonding, and thereby the β-sheet arrangement, remains stable over the remainder of the trajectory.

To probe local flexibility, we computed residue-resolved RMSF profiles for each protofibril (Fig. 2d). The two protofibrils that unravel at their C-termini correspond to sequence regions depleted in hydrophobic and aromatic residues (Fig. 2a,b), whereas the protofibril rich in such residues forms a stable, rigid segment. Electrostatic repulsion among the clustered lysines at Lys113 may further contribute to destabilizing the C-terminal regions, consistent with an abrupt drop in backbone hydrogen bond density in this region (Fig. 2e). In addition to these terminal effects, all protofibrils display a local peak in RMSF near Gly98, accompanied by a reduction in backbone hydrogen bonding. This feature likely reflects the intrinsic flexibility imparted by the glycine residue and does not compromise the integrity of the fibril core.

### Seeded fibril-growth simulations of Nup98 FG domains

To investigate how disordered FG domains interact with and grow upon pre-formed amyloid templates, we carried out seeded fibril-growth simulations for two Nup98 fibril polymorphs: Nup98_FG_(85–124) PM1 and Nup98_FG_(298–327). These two structures were selected because they exhibited the highest stability in our fibril-stability tests. For each system, we constructed a 30-layer fibril segment, which provides a sufficiently long and stable cross-β architecture to support growth at the fibril ends.

Since the simulation boxes are relatively small, monomers that bind to the fibril rapidly deplete the concentration of free chains in solution. To maintain a constant supply of unbound molecules, and thus a controlled driving force for elongation, we implemented a monomer-replenishment protocol [38]. Every 100 ns, the simulation was paused and all chains were classified as either free or associated with the fibril (either directly bound to the template or interacting with already bound chains). Additional monomers were then added to restore the concentration of free molecules to 1mM. This approach ensures that fibril growth is not limited by monomer depletion and allows us to monitor elongation, binding modes, and secondary nucleation events under steady solution conditions.

To illustrate the fibril-growth behavior observed in these simulations, we focus on the Nup98_FG_(85–124) PM1 system (Fig. 3). A representative snapshot after 10 μs shows extensive association of disordered monomers with the fibril seed, which retains its three-protofibril architecture. Newly attached chains accumulate along the fibril surface and gradually adopt ordered secondary structure, becoming incorporated into the growing assembly. Two distinct growth mechanisms can be identified from the zoom-in views. First, incoming monomers elongate the fibril by aligning with the existing β-sheet registry in a sequence-matched manner, thereby extending the ordered core with precise structural registration. Second, the fibril surface serves as a template for secondary nucleation, where surface-bound chains reorganize into β-sheet-rich structures that form additional protofibrillar assemblies branching from the original seed.

**Fig. 3.**
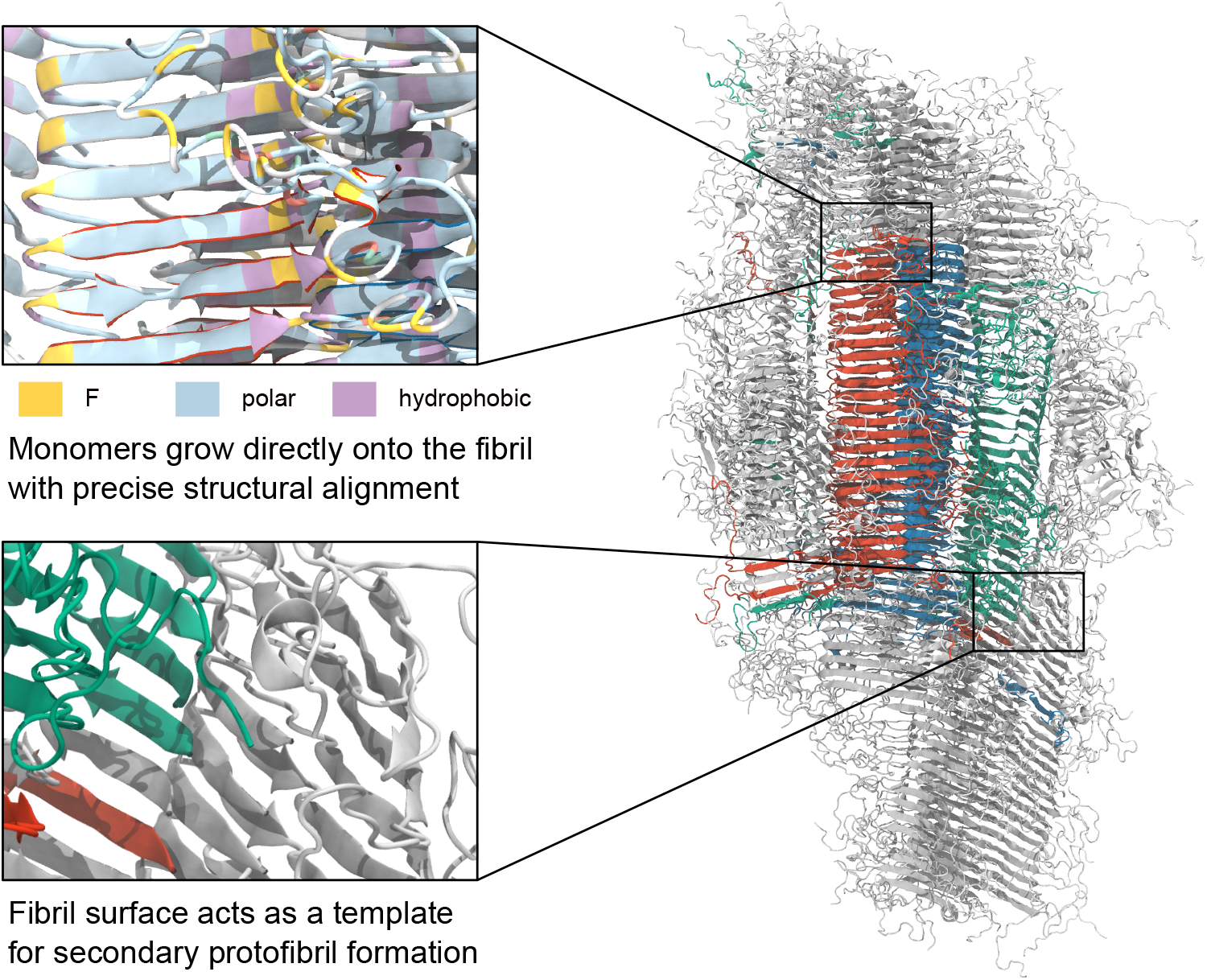
Seeded fibril-growth simulations of Nup98^FG^(85–124) PM1. Representative snapshot from a 10 µs seeded growth simulation of Nup98^FG^(85–124) PM1 (PDB ID: 7Q64), shown in secondary structure representation. The original fibril seed is colored blue, red, and green to distinguish the three protofibrils of the structure, while newly attached monomers are shown in gray. Zoom-in views highlight two key observations: (i) incoming monomers elongate the fibril through precise structural alignment with the existing β-sheet architecture, and (ii) the fibril surface serves as a template for the nucleation of secondary protofibrils.

Across independent replicas, we observe variation in the relative contribution of these mechanisms. In some simulations, fibril extension at the tips is more pronounced, whereas in others the formation of surface-associated condensates and secondary protofibrils dominates. Representative snapshots of additional replicas, as well as corresponding simulations of Nup98_FG_(298–327), are provided in Suppl. Figs. S8 and S9.

### Phase separation of FG-Nups using the 2BPA-HB model

To investigate whether the 2BPA-HB model can reproduce the phase separation behavior of FG-Nup domains, we performed coarse-grained simulations of selected FG-Nups using the same sequences and procedure as in our previous work [10]. Simulations were initiated from pre-formed condensates, which were then equilibrated to test whether the FG domains could maintain a stable phase-separated state. This approach allowed the system to reach equilibrium more efficiently compared to starting from a dispersed configuration.

We tested phase separation for the FG domains of five representative FG-Nups that together capture the range of FG domain behaviors observed across the NPC [10, 13]: Nup98, Nup100, Nup49, Nup159, and Nsp1. For Nsp1, we additionally simulated only the collapsed domain (residues 1–186), as the full FG domain exhibits a bimodal charge distribution that gives rise to a cohesive N-terminal domain and a noncohesive C-terminal domain [10]. For the GLFG-Nups (Nup98, Nup100, and Nup49) as well as the collapsed N-terminal domain of Nsp1, we observe phase separation into stable condensates (Fig. 4a), consistent with experiments [13, 14, 49] and our earlier results using the 1BPA model [10]. A notable difference, however, is that the dilute phases in the 2BPA-HB simulations are substantially less populated than in the 1BPA simulations, indicating stronger overall cohesiveness in the new model. As expected, Nup159 does not undergo phase separation (Fig. 4a), again in agreement with previous findings.

**Fig. 4.**
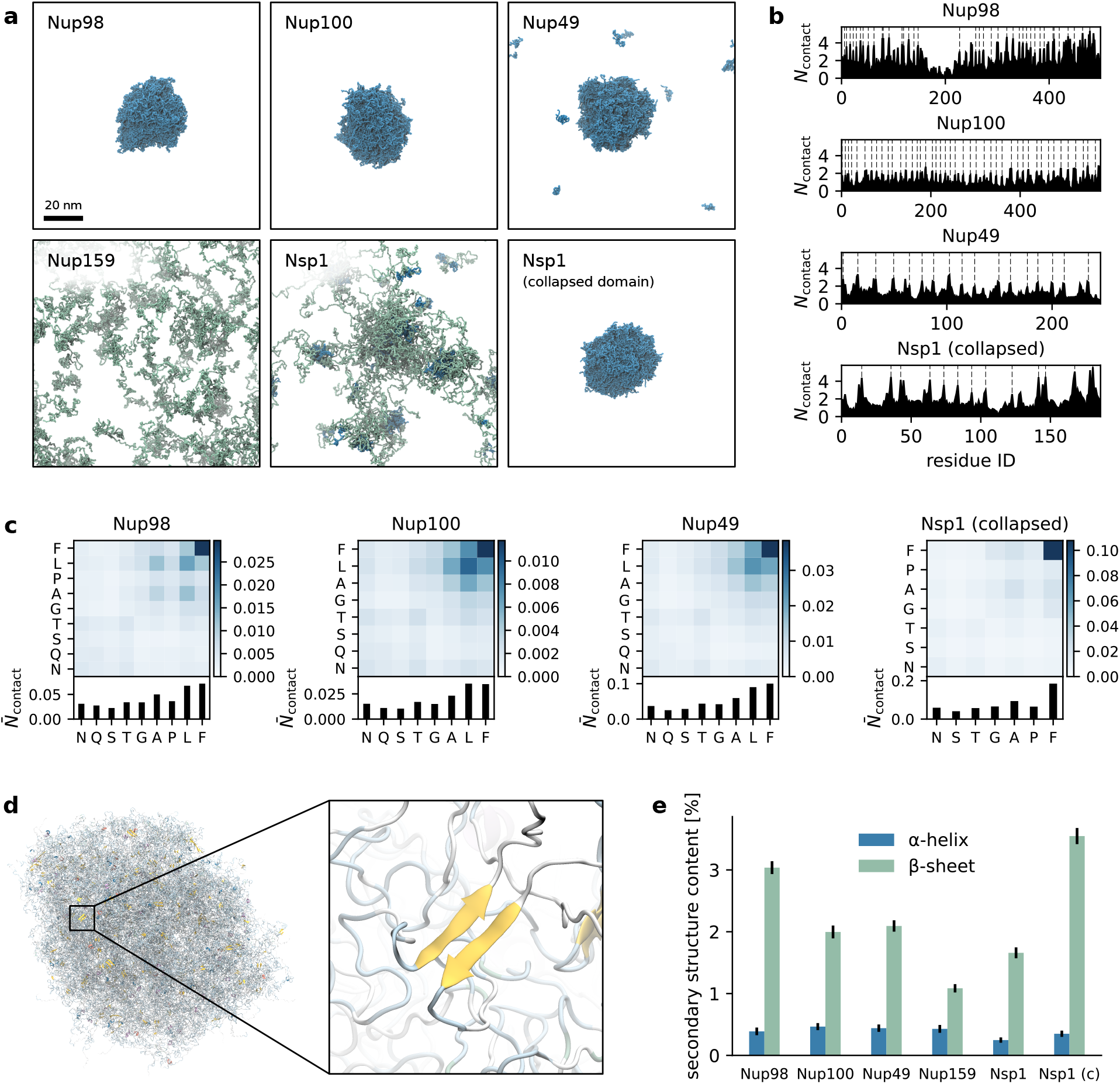
Phase separation behavior of selected FG-Nups in the 2BPA-HB model. (**a**) Representative simulation snapshots of the equilibrium state for each FG-Nup system. Cohesive domains are shown in blue and noncohesive domains in green. (**b**) Time-averaged number of intermolecular contacts per protein replica, resolved by residue index, for Nup98, Nup100, Nup49, and the collapsed domain of Nsp1. Dashed lines indicate the location of FG motifs. Full contact maps showing the intermolecular contacts per residue pair are provided in Suppl. Figs. S10 and S11. (**c**) Time-averaged intermolecular contacts per protein replica, resolved by residue type and normalized by residue abundance. Top panels display contact frequencies between all pairs of residue types; bottom panels show the total normalized contacts per residue type. Only residue types contributing at least 4% to the sequence are included. The corresponding non-normalized contact maps are shown in Suppl. Figs. S10 and S11. (**d**) Snapshot of the Nsp1 collapsed domain condensate backmapped to atomistic resolution and rendered in secondary-structure representation. The inset highlights a locally ordered β-sheet motif formed within the condensate. (**e**) Average secondary structure content of the FG-Nup condensates, showing the fraction of residues adopting α-helical or β-strand conformations. Percentages are computed over all residues and averaged over the equilibrium portion of the trajectories.

Interestingly, the full FG domain of Nsp1 does not form a single uniform condensate, yet exhibits cluster formation. These clusters are interacting primarily through the cohesive, collapsed N-terminal domain (which strongly phase separates on its own) and organize into smaller micelle-like structures. These micellar clusters remain interconnected over longer timescales, potentially stabilized by FG cross-links formed by the extended, noncohesive C-terminal domain.

To identify which residues contribute most to phase separation in these FG-Nups, we analyzed in-termolecular contact statistics within the condensates. We computed the time-averaged number of intermolecular contacts per protein replica for each residue in the sequence for Nup98, Nup100, Nup49, and the collapsed domain of Nsp1 (Fig. 4b). In all cases, contacts were most frequent at FG motifs, broadly consistent with trends observed previously in 1BPA simulations [10]. When contacts are grouped by residue type (Fig. 4c), phenylalanine residues consistently show the highest number of interactions across all systems. Compared to the 1BPA simulations, polar residues contribute more to phase separation, although the overall impact of the increased attractiveness between polar side chains (Eq. 5) remains modest.

Given the explicit backbone hydrogen bonding interactions in the 2BPA-HB model, we next examined whether FG-Nup condensates exhibit any spontaneous secondary structure formation. Simulation snapshots were backmapped to atomistic resolution and analyzed for secondary structure content (see Supplementary Methods for details). This analysis revealed that secondary structure elements are consistently present in all condensates. An illustrative example is shown in Fig. 4d, where the backmapped condensate of the Nsp1 collapsed domain exhibits transient β-sheet motifs. Quantitative analysis over the final 5 μs of each trajectory shows a small fraction of α-helical content (∼0.5%) and a more substantial β-sheet content (1% to 4%) across each of the FG-Nup systems (Fig. 4e). This level of secondary structure formation is significantly higher than observed in our earlier phase separation simulations using the 1BPA model (Suppl. Fig. S12), which lacks explicit backbone hydrogen bonding interactions. Despite the continuous presence of β-sheet motifs, these structures are highly dynamic and short-lived, and do not develop into stable, amyloid-like assemblies within the simulated timescales.

## Discussion

In this work, we set out to develop a CGMD model capable of capturing both the disordered and amyloid states of FG-Nups in a sequence-resolved manner. Building on our earlier 2BPA-Q framework [38], we introduced the 2BPA-HB model with the aim of constructing a single computational framework that can represent intrinsically disordered FG-Nups in solution, in liquid-like condensates, and in amyloid-like assemblies using the same residue-level description. By deriving side chain placement parameters from all-atom simulations and introducing explicit backbone hydrogen bonding interactions, the 2BPA-HB model incorporates sufficient structural detail to capture sequence-specific interactions, secondary structure formation, and the conformational preferences characteristic of intrinsically disordered proteins. At the same time, its coarse-grained resolution enables microsecond-scale simulations of systems far beyond the reach of all-atom simulations, allowing us to probe both the structural and phase separation behavior of FG domains under biologically relevant conditions.

Our simulations demonstrate that the 2BPA-HB model can maintain the structural stability of experimentally determined Nup98 fibril polymorphs across microsecond timescales. Polymorph 1 (PM1), in particular, preserves its characteristic three-protofibril architecture and β-sheet registry, with only modest rearrangements in local geometry. For other polymorphs, however, fibril stability depends on fibril length: we observed dissociation and re-association of monomers at the fibril ends. This behavior suggests that minimal fibril length (or equivalently a minimal local monomer concentration) is required to create and maintain a stable β-sheet core. Notably, this concentration-dependent stability is consistent with experiment, as the resolved Nup98 fibril structures were formed under high concentration conditions [20, 21]. Beyond global preservation of the fibril architecture, the model also displays sequence-dependent variations in local flexibility, including the increased mobility of C-terminal segments and the stabilizing roles of hydrophobic and aromatic residues within the fibril core. Seeded growth simulations further demonstrate that disordered FG fragments can align with the fibril template in a sequence-specific manner, adopt β-sheet conformations, and, in some cases, nucleating additional protofibrils via secondary growth.

We applied the 2BPA-HB model to study the phase separation behavior of the human Nup98 and several yeast FG-Nups that we have previously studied using the 1BPA model. Consistent with earlier results, the FG domains of Nup98, Nup100, Nup49, and the cohesive N-terminal region of Nsp1 all formed stable condensates. In all cases, FG motifs provide the dominant cohesive interactions, while polar or charged regions contribute more weakly, in agreement with experimental observations and previous modeling [8, 10, 50]. With the 2BPA-HB model’s refined distribution of attractive forces, such as enhanced specificity of F–F (π–π) interactions and increased attraction between polar side chains, these interaction patterns emerge naturally in residue-resolved contact maps. Moreover, the higher structural resolution of our model reveals transient β-structure formation within liquid-like condensates, offering new insight into the coexistence of disorder and local structural order in FG domains. This dual capability, capturing both phase separation and fibril formation within a single sequence-resolved framework, distinguishes the 2BPA-HB model from other models and opens new possibilities for studying the interplay between cohesiveness, conformational heterogeneity, and structured aggregation of intrinsically disordered proteins. Notably, we do not observe spontaneous formation of higher-order amyloid assemblies or fibril nucleation events within the simulated condensates. Although transient β-sheet motifs continuously form and dissolve, they do not mature into stable cross-β architectures on the microsecond timescales accessible here. Primary nucleation is a stochastic process and therefore unlikely to be captured within the microsecond timescales sampled in our simulations [51].

A key design choice in the 2BPA-HB model is the use of a residue-specific interaction center rather than the center of mass of the side chain. This approach significantly changes the side chain placement for bulky, asymmetric, or aromatic residues and is essential for maintaining stable Nup98 fibrils with only small deviations from the experimental structures. At the same time, the model assigns each side chain a fixed position relative to the backbone, even though all-atom side chains are dynamic and occupy multiple rotameric states (Suppl. Fig. S1). By selecting the most probable interaction center position for each residue type, we capture the dominant configuration in a way that works well in most environments. Nevertheless, in some fibril snapshots we observe cases where the fixed interaction site does not coincide with the side chain’s expected rotamer in the experimental structure, leading to suboptimal packing or an underestimation of stabilizing contacts (Suppl. Fig. S13). A coarse-grained representation that allows dynamic sampling of side chain conformers, as implemented in models such as UNRES [52] and HyRes [37, 53], would naturally address this limitation and may further improve the accuracy of β-sheet packing and interstrand interactions.

Several coarse-grained strategies have been developed to study intrinsically disordered proteins, but none simultaneously capture sequence-resolved phase separation and amyloid-like structure within a single framework. Models such as HPS [28], Mpipi [30], and related residue-level potentials have proven highly successful for predicting LLPS behavior, but they lack explicit backbone directionality and therefore cannot form stable β-sheets. At the other end of the spectrum, structure-focused CGMD models such as AWSEM [54] and UNRES [52] incorporate detailed secondary-structure energetics but are not parameterized for disordered chains. MARTINI [55] offers greater chemical detail but requires elastic restraints to maintain secondary structure, limiting its ability to model spontaneous β-sheet formation. An emerging alternative is CGSchNet [56], a machine-learned coarse-grained architecture that represents proteins using a small number of physically interpretable sites, but with the added flexibility of learning effective potentials directly from atomistic simulations. This makes CGSchNet a promising candidate for tackling the same class of problems addressed here, although large-scale simulations of multi-protein systems have not yet been demonstrated [57]. In this landscape, the 2BPA-HB model occupies a unique niche: it retains the efficiency and sequence specificity of residue-level models while incorporating explicit, directional hydrogen bond interactions, enabling the simultaneous description of disordered states, condensates, and fibrillar structures.

An interesting next step will be to unify the 2BPA-Q and 2BPA-HB frameworks into a single transferable coarse-grained model capable of handling both FG-Nups and low-complexity amyloid-forming domains. The geometry of the glutamine side chain is nearly identical in both models, suggesting that polyQ assemblies should in principle remain accessible within the 2BPA-HB representation. However, the nonbonded interaction parameters differ substantially: in its current parametrization, 2BPA-HB produces polyQ chains that are considerably more extended than those obtained with 2BPA-Q, which was explicitly calibrated against all-atom simulations of polyQ monomers [38]. Achieving full compatibility between the two models would therefore require a systematic reparametrization of the nonbonded interactions, such that a unified model can simultaneously reproduce polyQ monomer scaling behavior, hydrodynamic radii of FG-Nups, and the ability to sustain stable fibrils formed by both polyQ and FG domains.

A second major direction is the simulation of nuclear pore complexes, where the transient formation of secondary structure may influence the conformational and transport properties of the central channel [22, 58]. With the 2BPA-HB model, it becomes feasible to probe how local β-structure formation alters barrier permeability, and how this dynamic barrier is modulated by nuclear transport receptors and Nsp1 to remain functional [13, 18, 19, 23]. Finally, recent work has raised the question of how neurodegeneration-linked proteins, such as Tau or TDP-43, perturb FG-Nup organization when they aberrantly accumulate within the pore [59–61]. Because the 2BPA-HB model captures both disordered and amyloid-like conformational states within a single framework, it provides a promising platform for exploring how such aggregates reshape FG-Nup dynamics and, more broadly, how nucleocytoplasmic transport defects may arise from pathological protein aggregation.

Taken together, our work establishes the 2BPA-HB model as a versatile coarse-grained framework that unifies two critical aspects of FG-Nup behavior: their ability to form dynamic, liquid-like assemblies and their capacity to adopt structured, β-sheet-rich amyloid conformations under appropriate conditions. By bridging these two regimes within a single representation, the model creates exciting new opportunities for multiscale studies of nucleocytoplasmic transport, fibril assembly, and pathological aggregation that were previously beyond reach. More broadly, by combining sequence resolution with explicit secondary-structure formation at low computational cost, the 2BPA-HB model provides a foundation for exploring how disorder, cohesiveness, and structure collectively shape the behavior of low-complexity domains in health and disease.

## Acknowledgements

This work was financially supported by the Netherlands Organization of Scientific Research grant no. OCENW.GROOT.2019.068. This work made use of the Dutch national e-infrastructure with the support of the SURF Cooperative using grant no. EINF-9807. We thank the Center for Information Technology of the University of Groningen for their support and for providing access to the Hábrók high performance computing cluster.

## Supplementary methods : Simulation systems and setup

### Coarse-grained simulations

All CGMD simulations were performed using the 2BPA-HB model and carried out with the GRO-MACS [47] simulation package (version 2019.6). A modified implementation of the 3OUT virtual site scheme was used to construct the backbone hydrogen bonding beads required by the model, following the procedure described in Ref. [38]. Unless stated otherwise, simulations were conducted at a temperature of 300 K using Langevin dynamics with a time step of 0.02 ps and an inverse friction coefficient of *γ*^−1^ = 2 ps.

#### Fibril equilibration simulation

Fibril equilibration simulations were performed using experimentally resolved fibril structures. Starting from the reported atomic coordinates obtained from the corresponding PDB files, each structure was extended along the fibril axis to construct fibers consisting of 20, 30, or 50 layers. Here, a layer is defined as one β-strand contributed by each protofibril within the fibril cross-section. The resulting fibril segments were placed in periodic simulation boxes sufficiently large to prevent interactions between periodic images and were simulated for 5 μs to assess the stability of the fibrillar architecture.

#### Seeded fibril growth simulations

Seeded growth simulations were initialized from preformed 30-layer fibril segments placed in a periodic simulation box with side length 70 nm, together with disordered FG fragments randomly distributed in solution. The number of free monomers was chosen to yield an initial concentration of 1mM. As monomers attach to the fibril, the concentration of the dilute phase decreases over time. To maintain a constant driving force for fibril elongation, simulations were paused every 100 ns and the size of the largest molecular cluster was determined using the `gmx clustsize` utility with a distance cutoff of 0.55 nm. Chains belonging to this cluster, including both ordered fibrillar chains and disordered molecules in contact with the fibril, were excluded when estimating the dilute phase concentration. Additional disordered monomers were then inserted at random positions and orientations to restore the free monomer concentration to its target value, after which the simulation was resumed.

#### Phase separation simulations

Phase separation simulations of FG-Nups were performed following the same general protocol as in our previous work using the 1BPA model [10]. Briefly, initial configurations were generated by first equilibrating a dense monomer solution in the NVT ensemble to promote mixing, followed by an isotropic expansion of the simulation box to reach a target concentration of approximately 7mg/mL, yielding an preformed condensate. This procedure significantly accelerates equilibration compared to starting from a homogeneous dilute solution. System sizes were chosen such that the total number of amino acids was sufficiently large to minimize finite-size effects. Each system was simulated for 10 μs, with dynamic equilibrium typically reached within the first 5 μs (Suppl. Fig. S14).

### All-atom simulations

All-atom MD simulations were performed using the GROMACS [47] simulation package (version 2021.3) with the amber99SB-disp force field [62] and the TIP4P-D water model. Equations of motion were integrated using the leap-frog algorithm with a 2 fs time step, and all bonds involving hydrogen atoms were constrained using the LINCS algorithm [63]. Short-range van der Waals interactions were truncated at 1.0 nm, and long-range electrostatics were treated using the particle mesh Ewald method [64] with a real-space cutoff of 1.2 nm, a Fourier spacing of 0.125 nm, and fourth-order interpolation. Long-range dispersion corrections were applied to energy and pressure. Temperature was maintained at 300K using the stochastic velocity-rescaling thermostat [65] (coupling constant 1.0 ps), and pressure was controlled at 1 bar using the Parrinello–Rahman barostat [66] (coupling constant 2.0 ps, compressibility 4.5 × 10^−5^ bar^−1^).

Simulations were performed for five FG-Nup fragments (Nup98_FG_(85–124), Nup98_FG_(165–204), Nsp1, Nup145, and Nup60; see Suppl. Table S2 for details). Each fragment was placed in an extended conformation in a periodic simulation box, solvated with explicit water, and neutralized with sodium and chloride ions to an ionic strength of 100mM. Following energy minimization, systems were equilibrated for 1 ns in the NVT ensemble and subsequently for 1 ns in the NPT ensemble. Production simulations were then performed in the NPT ensemble for at least 500 ns. The resulting trajectories were used to extract side chain conformational statistics for coarse-grained model parameterization.

## Supplementary methods: Analysis of coarse-grained simulations

### Intermolecular contact analysis

For phase separated systems, only interactions between FG-Nup molecules belonging to the condensed phase were considered. For each trajectory frame, molecules in the condensate were identified using the `gmx clustsize` utility of GROMACS, defining the largest molecular cluster as the condensate. A protein chain was classified as part of the condensate if at least one of its beads was within 0.55 nm of any bead belonging to the largest cluster.

For each frame, intermolecular residue–residue contact matrices were constructed by considering all unique pairs of FG-Nup chains within the condensate. A residue was defined as being in contact with another residue if either the backbone (BB) or side chain (SC) bead of the first residue was within 0.55 nm of the BB or SC bead of the second residue. For a system containing *N*_Nup_ molecules in the condensate, this procedure yields *N*_Nup_(*N*_Nup_ − 1)/2 intermolecular contact matrices per frame. The average intermolecular contact map was obtained by summing all pairwise contact matrices within a frame and normalizing by the number of molecules in the condensate. These contact maps were subsequently averaged over all analyzed frames to yield the final contact probabilities.

### Secondary structure analysis

Secondary structure content was quantified by backmapping coarse-grained simulation frames to atomistic resolution followed by standard secondary structure assignment. For each selected simulation frame, the coarse-grained configuration was converted to an all-atom rep-resentation using CGBack [67], a Python-based backmapping tool that employs a diffusion-based generative model to reconstruct atomistic protein structures from coarse-grained coordinates. CGBack uses the positions of the backbone (BB) beads as input and outputs a complete all-atom model, and is capable of handling large-scale systems by backmapping all molecules in a given frame simultaneously. Although CGBack provides refinement options to resolve steric overlaps and optimize local geometry, applying such procedures to the full phase separation systems (containing ∼7 × 10^4^ residues and hundreds of analyzed frames) would be computationally prohibitive. We therefore performed backmapping without additional refinement or energy minimization. The resulting atomistic structures were visually inspected and found to be sufficiently accurate for secondary structure classification.

Secondary structure assignment was performed using STRIDE [68], which identifies secondary structural elements based on hydrogen bonding patterns and backbone geometry derived from atomic coordinates. Residues assigned to extended (E) or β-bridge (B) conformations were classified as β-sheet, while residues assigned as helix (H) were classified as α-helical. The fraction of residues in α or β conformations was calculated by dividing the number of residues in each class by the total number of residues in the system. For each simulation, secondary structure content was averaged over the final 5 μs of the trajectory, with frames sampled every 10 ns. While β-sheet conformations are robustly captured using this procedure, we note that helical conformations formed in the 2BPA-HB model are not always fully recognized by STRIDE. In particular, some helices deviate slightly from the geometric criteria used by STRIDE, leading to a potential underestimation of α-helical content.

## Supplementary Tables

**Suppl. Table S1.**
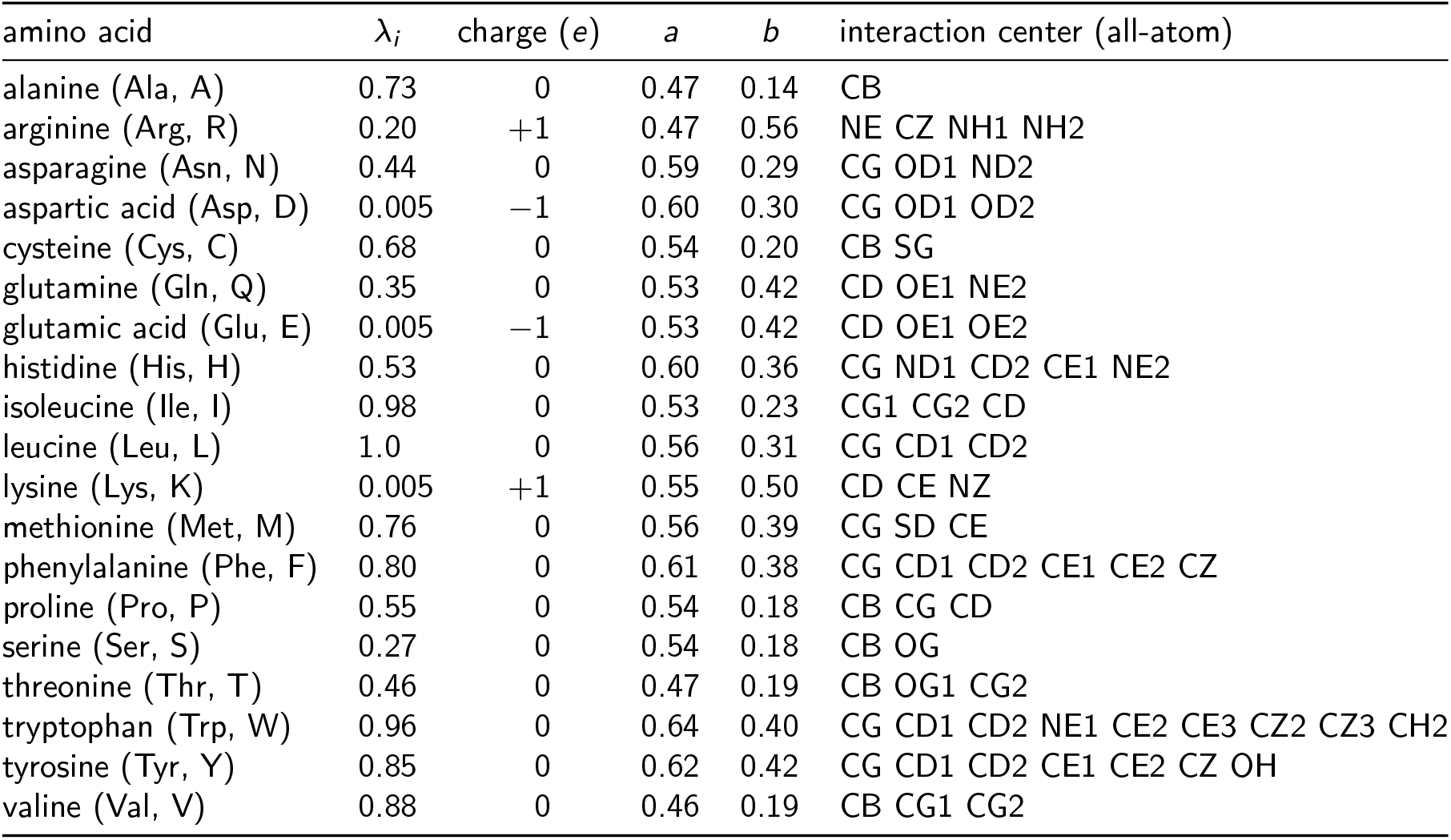
SC parameters for each amino acid in the 2BPA-HB model. The first column lists the amino acid name, followed by the relative hydrophobicity λ_*i*_ and net charge (*e*) of the side chain bead. Columns three and four show the optimal *a* and *b* parameters, determined from the most frequently occurring combination in the all-atom MD simulations. The final column lists the atoms used to define the side chain interaction center in the all-atom representation. For all amino acids, the backbone bead (BB) is identical and has a fixed λ_BB_ = 0.55.

| amino acid | $\lambda_i$ | charge ($e$) | $a$ | $b$ | interaction center (all-atom) |
| --- | --- | --- | --- | --- | --- |
| alanine (Ala, A) | 0.73 | 0 | 0.47 | 0.14 | CB |
| arginine (Arg, R) | 0.20 | +1 | 0.47 | 0.56 | NE CZ NH1 NH2 |
| asparagine (Asn, N) | 0.44 | 0 | 0.59 | 0.29 | CG OD1 ND2 |
| aspartic acid (Asp, D) | 0.005 | -1 | 0.60 | 0.30 | CG OD1 OD2 |
| cysteine (Cys, C) | 0.68 | 0 | 0.54 | 0.20 | CB SG |
| glutamine (Gln, Q) | 0.35 | 0 | 0.53 | 0.42 | CD OE1 NE2 |
| glutamic acid (Glu, E) | 0.005 | -1 | 0.53 | 0.42 | CD OE1 OE2 |
| histidine (His, H) | 0.53 | 0 | 0.60 | 0.36 | CG ND1 CD2 CE1 NE2 |
| isoleucine (Ile, I) | 0.98 | 0 | 0.53 | 0.23 | CG1 CG2 CD |
| leucine (Leu, L) | 1.0 | 0 | 0.56 | 0.31 | CG CD1 CD2 |
| lysine (Lys, K) | 0.005 | +1 | 0.55 | 0.50 | CD CE NZ |
| methionine (Met, M) | 0.76 | 0 | 0.56 | 0.39 | CG SD CE |
| phenylalanine (Phe, F) | 0.80 | 0 | 0.61 | 0.38 | CG CD1 CD2 CE1 CE2 CZ |
| proline (Pro, P) | 0.55 | 0 | 0.54 | 0.18 | CB CG CD |
| serine (Ser, S) | 0.27 | 0 | 0.54 | 0.18 | CB OG |
| threonine (Thr, T) | 0.46 | 0 | 0.47 | 0.19 | CB OG1 CG2 |
| tryptophan (Trp, W) | 0.96 | 0 | 0.64 | 0.40 | CG CD1 CD2 NE1 CE2 CE3 CZ2 CZ3 CH2 |
| tyrosine (Tyr, Y) | 0.85 | 0 | 0.62 | 0.42 | CG CD1 CD2 CE1 CE2 CZ OH |
| valine (Val, V) | 0.88 | 0 | 0.46 | 0.19 | CB CG1 CG2 |

**Suppl. Table S2.**
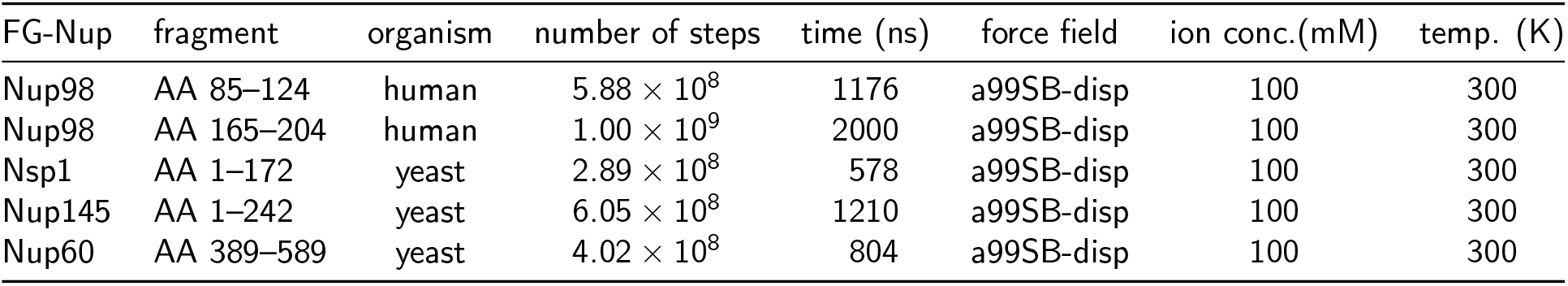
All-atom simulation details for FG-Nup fragments. Overview of the FG-Nup fragments simulated at atomistic resolution and the corresponding simulation parameters used to generate the trajectories for coarse-grained model parameterization.

| FG-Nup | fragment | organism | number of steps | time (ns) | force field | ion conc.(mM) | temp. (K) |
| --- | --- | --- | --- | --- | --- | --- | --- |
| Nup98 | AA 85–124 | human | $5.88 \times 10^8$ | 1176 | a99SB-disp | 100 | 300 |
| Nup98 | AA 165–204 | human | $1.00 \times 10^9$ | 2000 | a99SB-disp | 100 | 300 |
| Nsp1 | AA 1–172 | yeast | $2.89 \times 10^8$ | 578 | a99SB-disp | 100 | 300 |
| Nup145 | AA 1–242 | yeast | $6.05 \times 10^8$ | 1210 | a99SB-disp | 100 | 300 |
| Nup60 | AA 389–589 | yeast | $4.02 \times 10^8$ | 804 | a99SB-disp | 100 | 300 |

**Suppl. Table S3.** Predicted hydrodynamic radii for all Yamada FG-Nup segments in the 2BPA-HB model. All single-chain simulations are performed at *T* = 303K and salt concentration 150mM (*κ* = 1.27/nm). Hydrodynamic radii are calculated based on the BB beads only, using the HYDROPRO software [69]. Reported values are the averages obtained from 10 μs-long trajectories, sampling every 1 ns. Errors are calculated as |*R*_h;sim_ − *R*_h;exp_|*/R*_h;exp_.

| FG-Nup domain | AAs | length | $R_{h,\text{exp}}$ | $R_{h,\text{sim}}$ | error (%) |
| --- | --- | --- | --- | --- | --- |
| Nsp1n | 1–172 | 172 | 27.1 | 28.1 | 3.8 |
| Nup116m | 165–715 | 551 | 46.5 | 43.1 | 7.4 |
| Nup100n | 2–610 | 609 | 48.7 | 46.6 | 4.3 |
| Nup49 | 1–215 | 215 | 26.9 | 28.3 | 5.3 |
| Nup42 | 1–212 | 212 | 28.4 | 28.7 | 1.1 |
| Nup57 | 1–255 | 255 | 31.9 | 32.4 | 1.5 |
| Nup145N | 1–242 | 242 | 28.2 | 30.6 | 8.5 |
| Nup1c | 798–1076 | 279 | 32.4 | 31.2 | 3.7 |
| Nup159 | 441–881 | 441 | 55.4 | 56.7 | 2.3 |
| Nup60 | 389–539 | 151 | 31.3 | 32.3 | 3.1 |
| Nup1m | 220–797 | 578 | 67.9 | 67.3 | 0.8 |
| Nup2 | 186–561 | 376 | 59.8 | 53.1 | 11.2 |
| Nsp1m | 173–603 | 431 | 65.3 | 65.3 | 0.0 |
| Nup145Ns | 243–433 | 191 | 29.8 | 33.5 | 12.4 |
| Nup100s | 611–800 | 190 | 36.6 | 36.8 | 0.5 |
| Nup116s | 765–960 | 196 | 39.1 | 36.6 | 6.5 |
| **average** | – |  |  |  | **4.5** |

## Supplementary Figures

**Suppl. Fig. S1.**
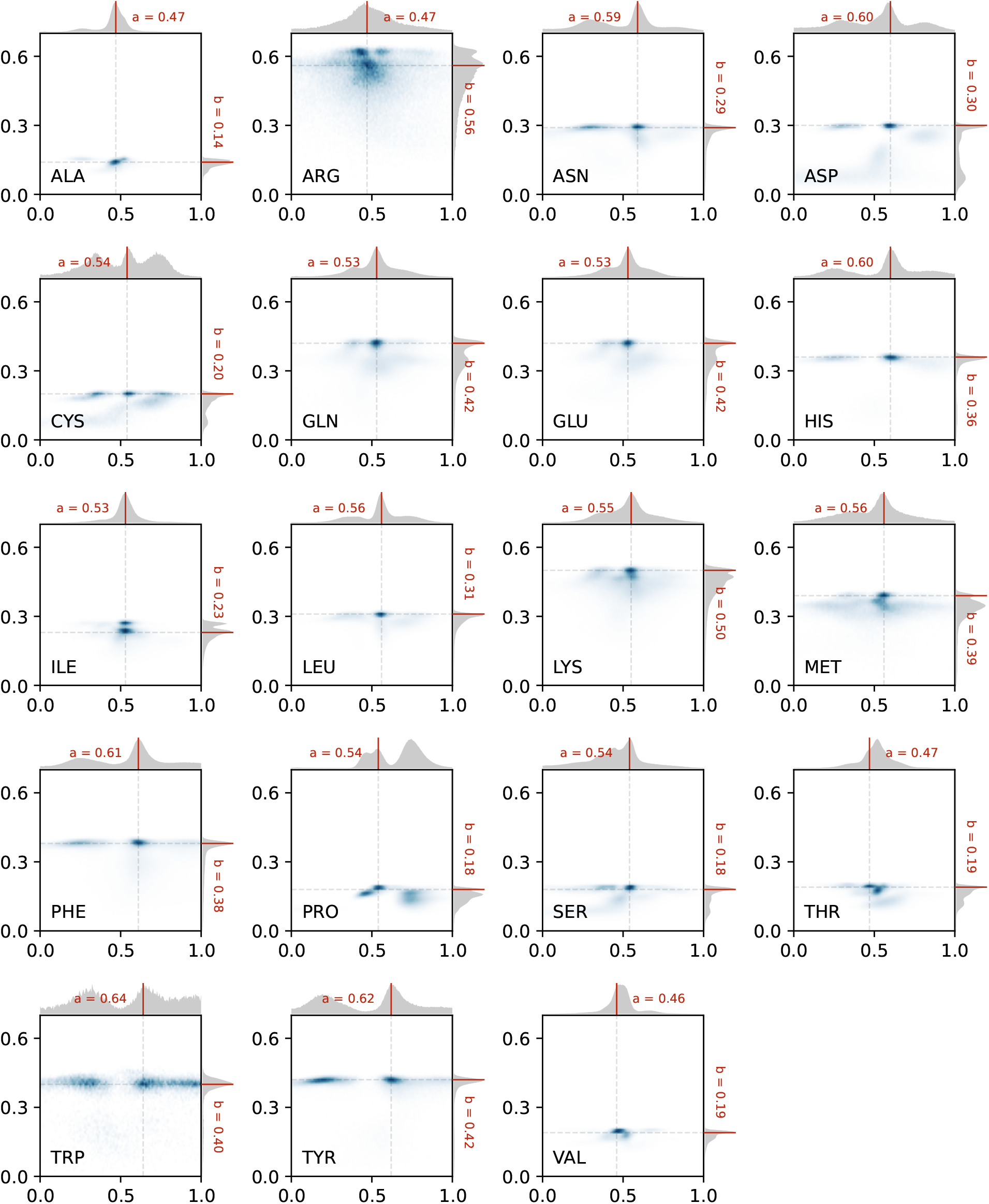
Two-dimensional distributions of side chain parameters (*a; b*) for all amino acid types. For each residue type, the two-dimensional histogram of side chain parameters (*a; b*) is shown, together with the corresponding one-dimensional marginal distributions along the top (*a*) and right (*b*) axes. Red lines indicate the (*a; b*) combination with the highest joint occurrence frequency (not the individually most frequent values), which were selected as the final side chain parameters in the 2BPA-HB model. These distributions were obtained from all-atom simulations of multiple FG-Nup segments (see Suppl. Table S2).

**Suppl. Fig. S2.**
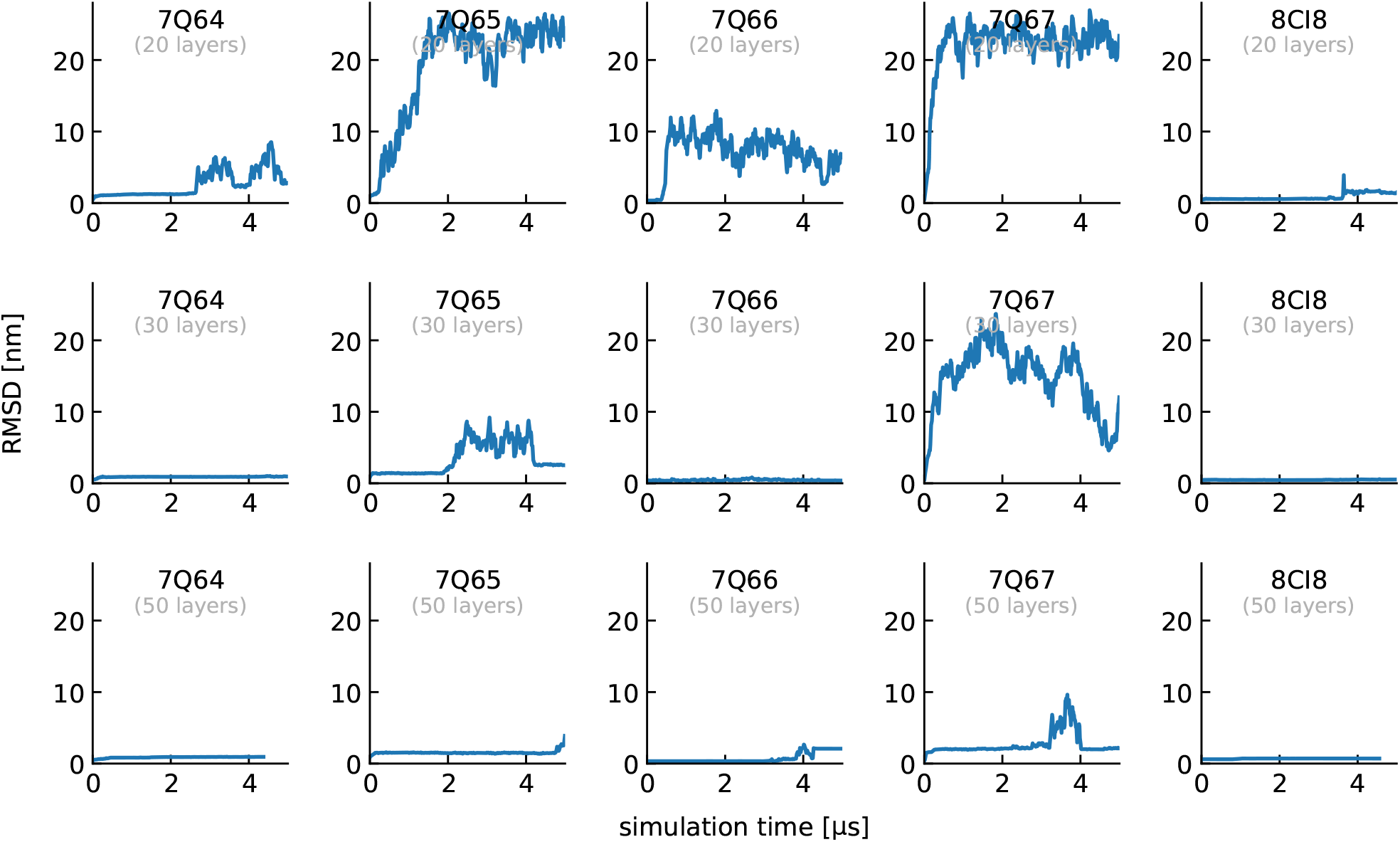
RMSD analysis of fibril stability for different systems and sizes. Root-mean-square deviation (RMSD) of fibril structures as a function of simulation time for systems with 20, 30, and 50 layers. Results are shown for multiple experimentally resolved Nup98 fibrils (PDB IDs: 7Q64, 7Q65, 7Q66, 7Q67, and 8CI8).

**Suppl. Fig. S3.**
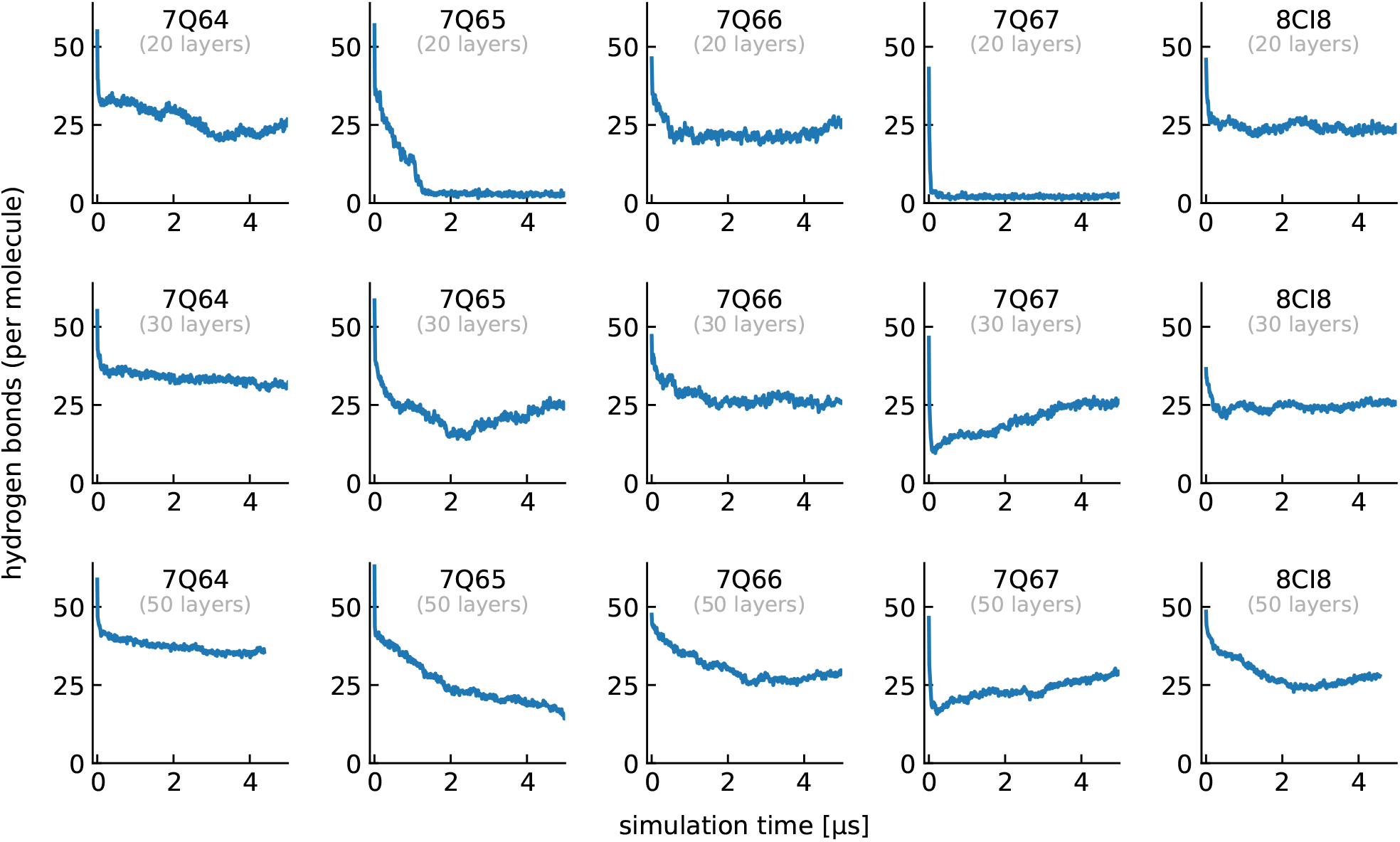
Hydrogen bond analysis of fibril stability for different systems and sizes. Average number of backbone hydrogen bonds per molecule as a function of simulation time for fibrils with 20, 30, and 50 layers. Results are shown for multiple experimentally resolved Nup98 fibrils (PDB IDs: 7Q64, 7Q65, 7Q66, 7Q67, and 8CI8).

**Suppl. Fig. S4.**
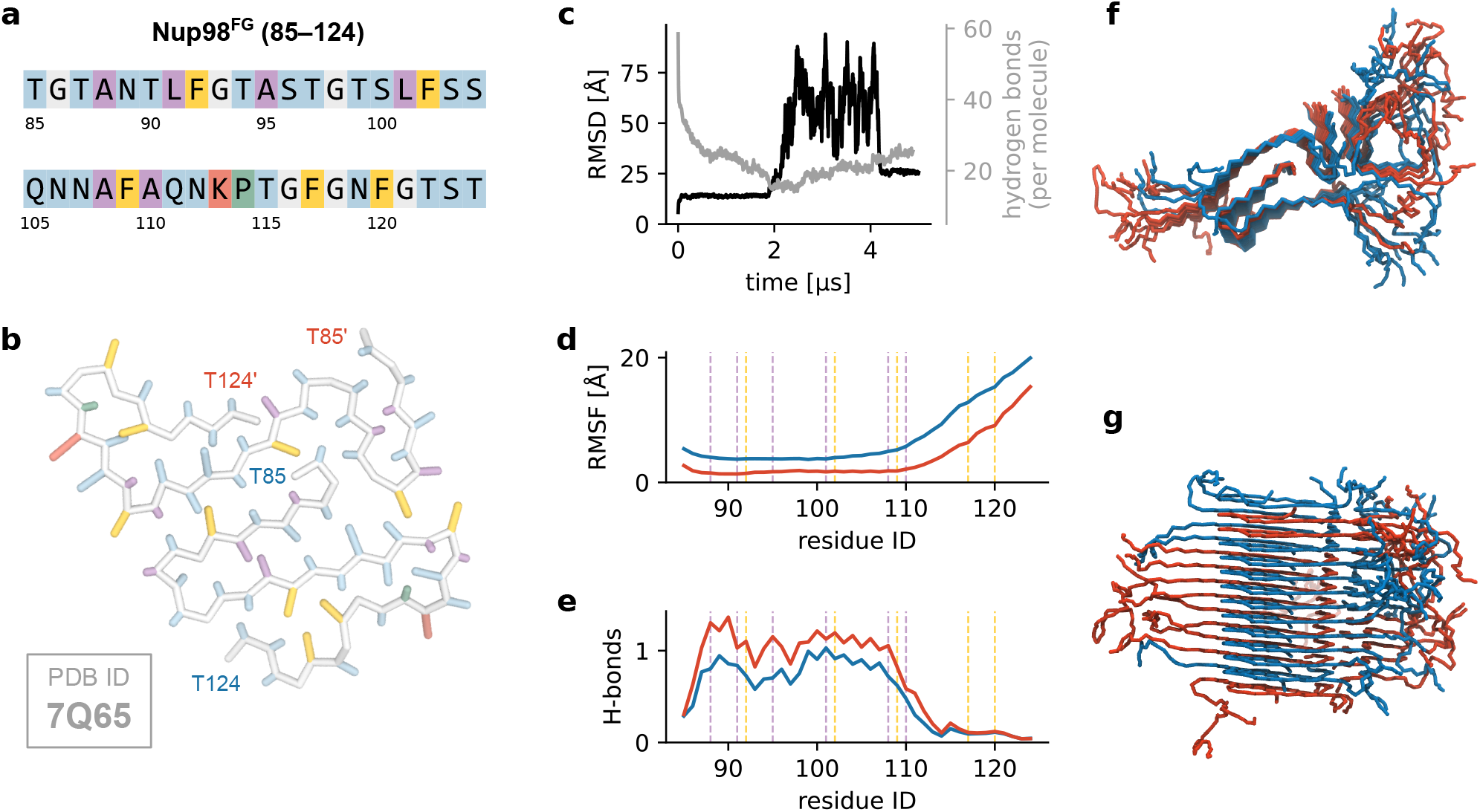
Stability and structural dynamics of the Nup98^FG^(85–124) PM3 fibril (PDB ID: 7Q65). **a** Amino acid sequence of Nup98^FG^(85–124), colored by residue type: polar (blue), hydrophobic (purple), aromatic (yellow), cationic (red), glycine (white), and proline (green). **b** Single cross-section of the fibril, highlighting the side chain packing within the coarse-grained representation and the resulting organization of the three protofibrils. **c** Time evolution of the RMSD of the fibril core (black) and the average number of backbone hydrogen bonds per molecule (gray). Only the middle 10 layers were used for the RMSD calculation, as terminal chains transiently dissociate from the fibril ends. Because each non-proline residue can donate and accept one hydrogen bond, the total number of hydrogen bonds can exceed the number of residues. **d** Average RMSF of residues in each protofibril, calculated from the middle 10 layers and color coded as in panels (**f**) and (**g**). Note that the protofibrils do not have the exact same length as in the resolved structure some terminal residues were missing. Dashed lines indicate the positions of phenylalanine (yellow) and hydrophobic (purple) residues. **e** Average number of backbone hydrogen bonds per residue for each protofibril. **f**,**g** Top (**f**) and side (**g**) views of the equilibrated 30-layer fibril at *t*\* = 5 μs, illustrating preservation of the overall protofibril architecture.

**Suppl. Fig. S5.**
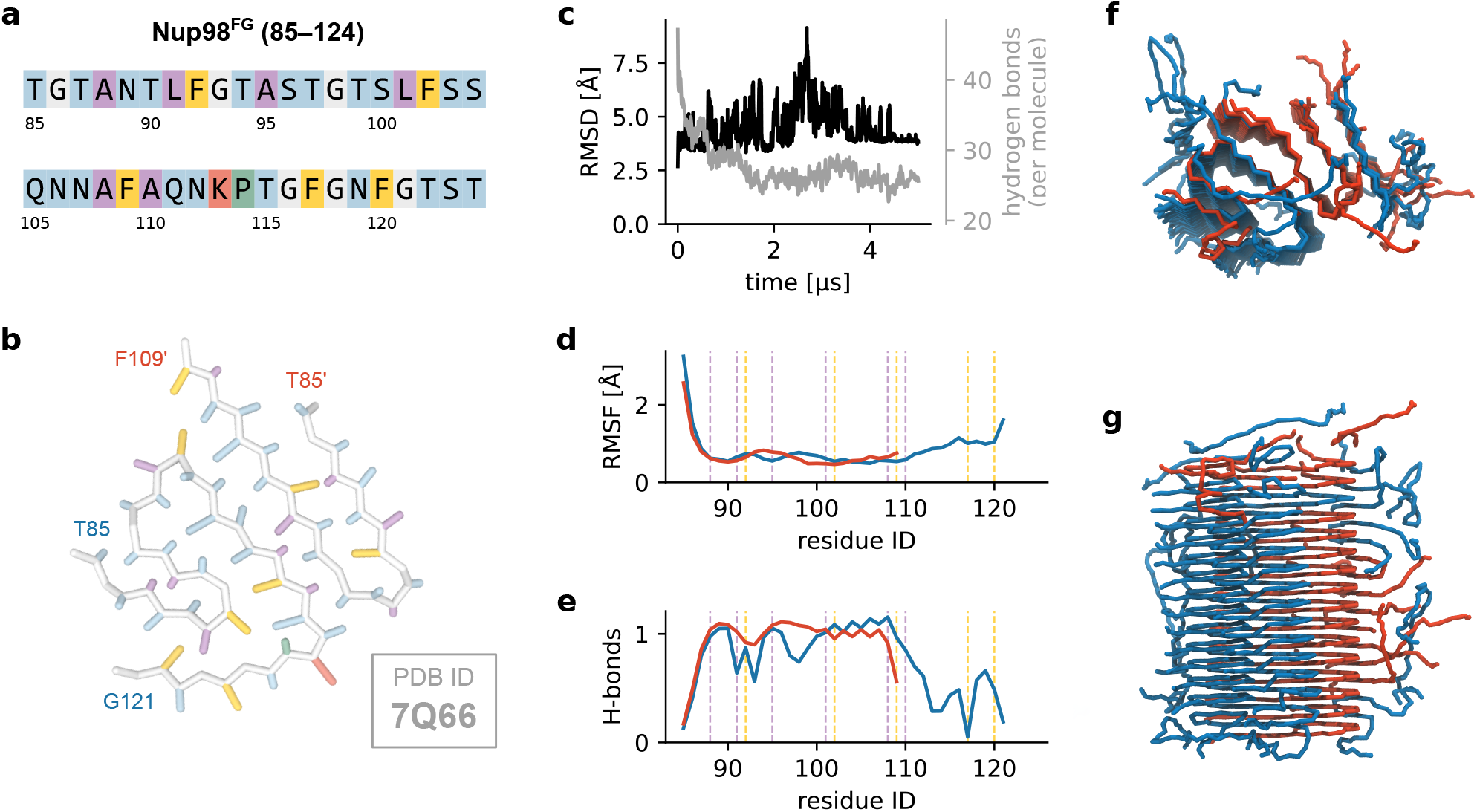
Stability and structural dynamics of the Nup98^FG^(85–124) PM3 fibril (PDB ID: 7Q66). **a** Amino acid sequence of Nup98^FG^(85–124), colored by residue type: polar (blue), hydrophobic (purple), aromatic (yellow), cationic (red), glycine (white), and proline (green). **b** Single cross-section of the fibril, highlighting the side chain packing within the coarse-grained representation and the resulting organization of the three protofibrils. **c** Time evolution of the RMSD of the fibril core (black) and the average number of backbone hydrogen bonds per molecule (gray). Only the middle 10 layers were used for the RMSD calculation, as terminal chains transiently dissociate from the fibril ends. Because each non-proline residue can donate and accept one hydrogen bond, the total number of hydrogen bonds can exceed the number of residues. **d** Average RMSF of residues in each protofibril, calculated from the middle 10 layers and color coded as in panels (**f**) and (**g**). Note that the protofibrils do not have the exact same length as in the resolved structure some terminal residues were missing. Dashed lines indicate the positions of phenylalanine (yellow) and hydrophobic (purple) residues. **e** Average number of backbone hydrogen bonds per residue for each protofibril. **f**,**g** Top (**f**) and side (**g**) views of the equilibrated 30-layer fibril at *t*\* = 5 μs, illustrating preservation of the overall protofibril architecture.

**Suppl. Fig. S6.**
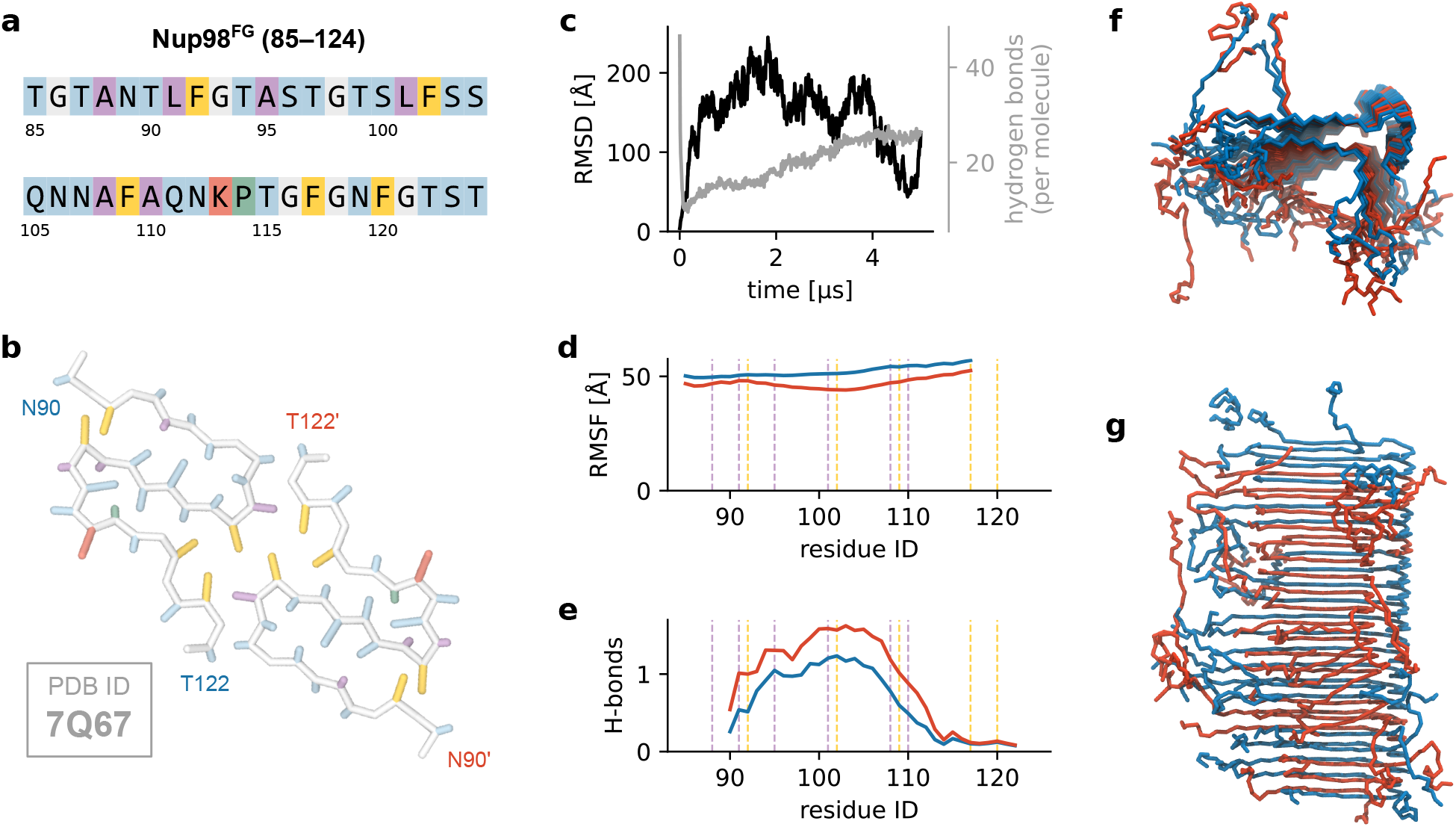
Stability and structural dynamics of the Nup98^FG^(85–124) PM4 fibril (PDB ID: 7Q67). **a** Amino acid sequence of Nup98^FG^(85–124), colored by residue type: polar (blue), hydrophobic (purple), aromatic (yellow), cationic (red), glycine (white), and proline (green). **b** Single cross-section of the fibril, highlighting the side chain packing within the coarse-grained representation and the resulting organization of the three protofibrils. **c** Time evolution of the RMSD of the fibril core (black) and the average number of backbone hydrogen bonds per molecule (gray). Only the middle 10 layers were used for the RMSD calculation, as terminal chains transiently dissociate from the fibril ends. Note that this fibril has refolded to a different structure. Because each non-proline residue can donate and accept one hydrogen bond, the total number of hydrogen bonds can exceed the number of residues. **d** Average RMSF of residues in each protofibril, calculated from the middle 10 layers and color coded as in panels (**f**) and (**g**). Note that the protofibrils do not have the exact same length as in the resolved structure some terminal residues were missing. Dashed lines indicate the positions of phenylalanine (yellow) and hydrophobic (purple) residues. **e** Average number of backbone hydrogen bonds per residue for each protofibril. **f**,**g** Top (**f**) and side (**g**) views of the equilibrated 30-layer fibril at *t*\* = 5 μs, illustrating preservation of the overall protofibril architecture.

**Suppl. Fig. S7.**
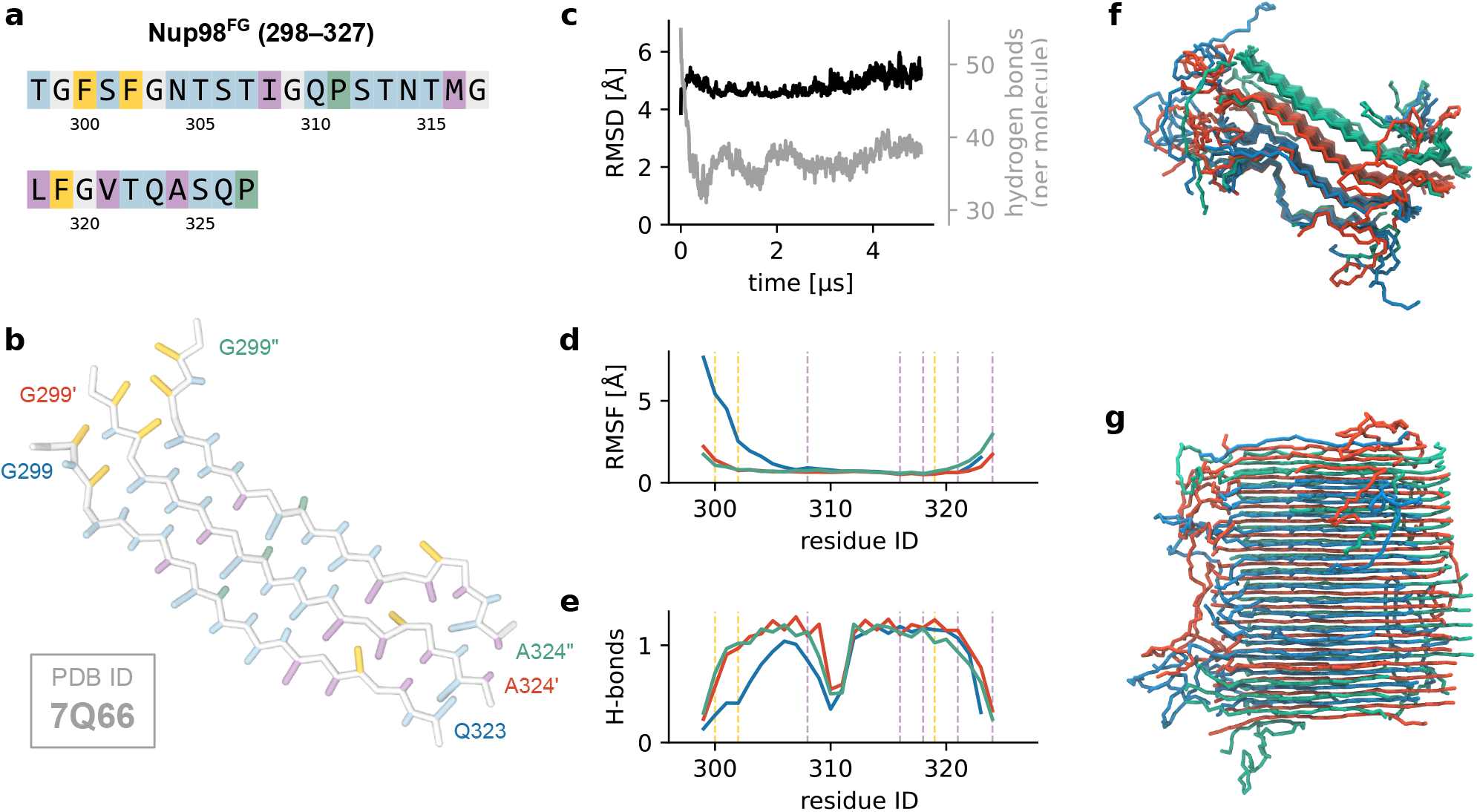
Stability and structural dynamics of the Nup98^FG^(298–327) fibril (PDB ID: 8CI8). **a** Amino acid sequence of Nup98^FG^(298–327), colored by residue type: polar (blue), hydrophobic (purple), aromatic (yellow), cationic (red), glycine (white), and proline (green). **b** Single cross-section of the fibril, highlighting the side chain packing within the coarse-grained representation and the resulting organization of the three protofibrils. **c** Time evolution of the RMSD of the fibril core (black) and the average number of backbone hydrogen bonds per molecule (gray). Only the middle 10 layers were used for the RMSD calculation, as terminal chains transiently dissociate from the fibril ends. Because each non-proline residue can donate and accept one hydrogen bond, the total number of hydrogen bonds can exceed the number of residues. **d** Average RMSF of residues in each protofibril, calculated from the middle 10 layers and color coded as in panels (**f**) and (**g**). Note that the protofibrils do not have the exact same length as in the resolved structure some terminal residues were missing. Dashed lines indicate the positions of phenylalanine (yellow) and hydrophobic (purple) residues. **e** Average number of backbone hydrogen bonds per residue for each protofibril. **f**,**g** Top (**f**) and side (**g**) views of the equilibrated 30-layer fibril at *t*\* = 5 μs, illustrating preservation of the overall protofibril architecture.

**Suppl. Fig. S8.**
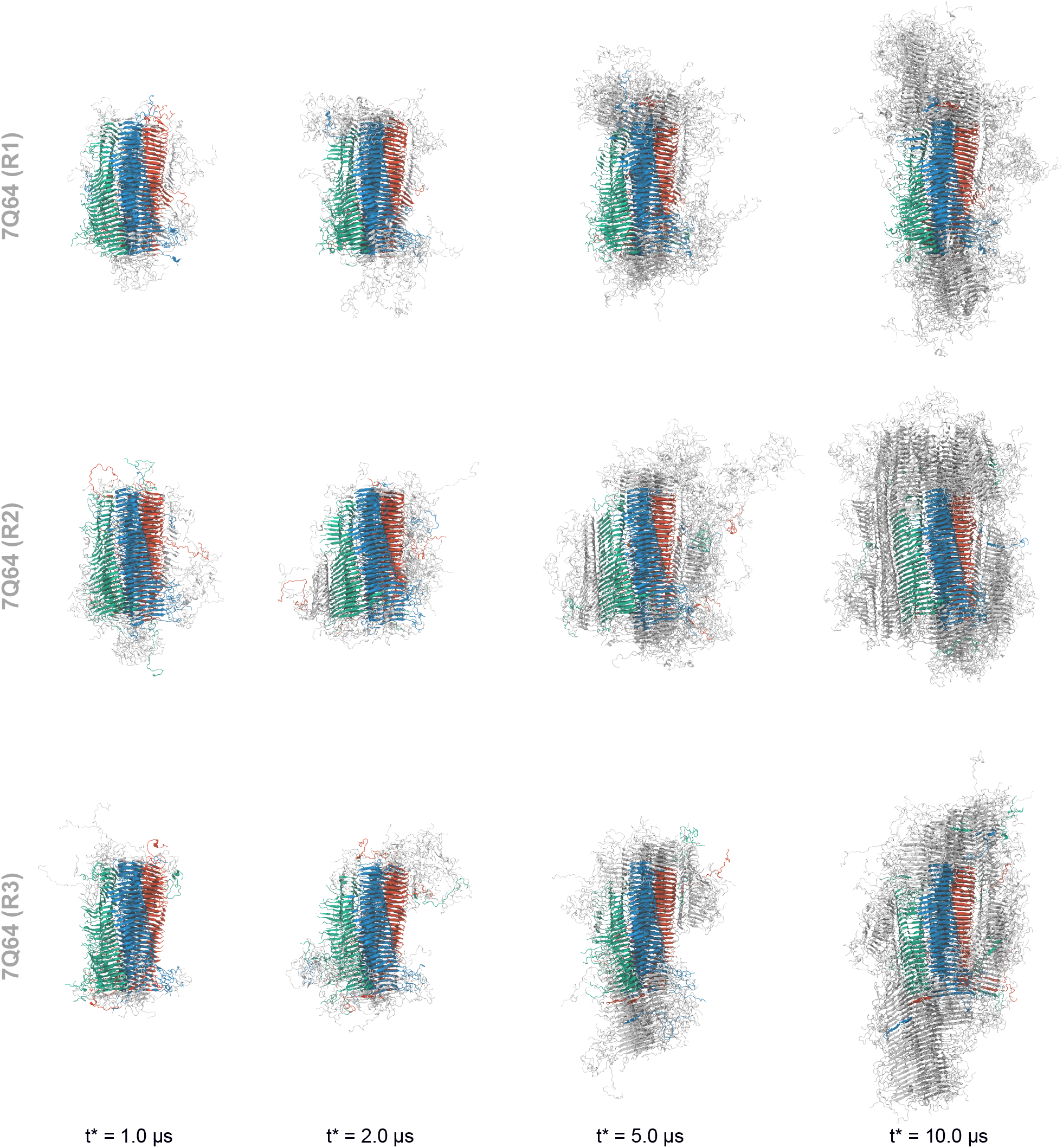
Representative snapshots of seeded fibril growth simulations of Nup98^FG^(85–124) (PM1). Snapshots from three independent simulations, each initialized from a 30-layer fibril seed of Nup98^FG^(85–124) (PDB ID: 7Q64), are shown at 1.0 μs, 2.0 μs, 5.0 μs, and 10.0 μs. The three protofibrils of the original seed are color-coded in red, green, and blue, while newly attached monomers are shown in gray.

**Suppl. Fig. S9.**
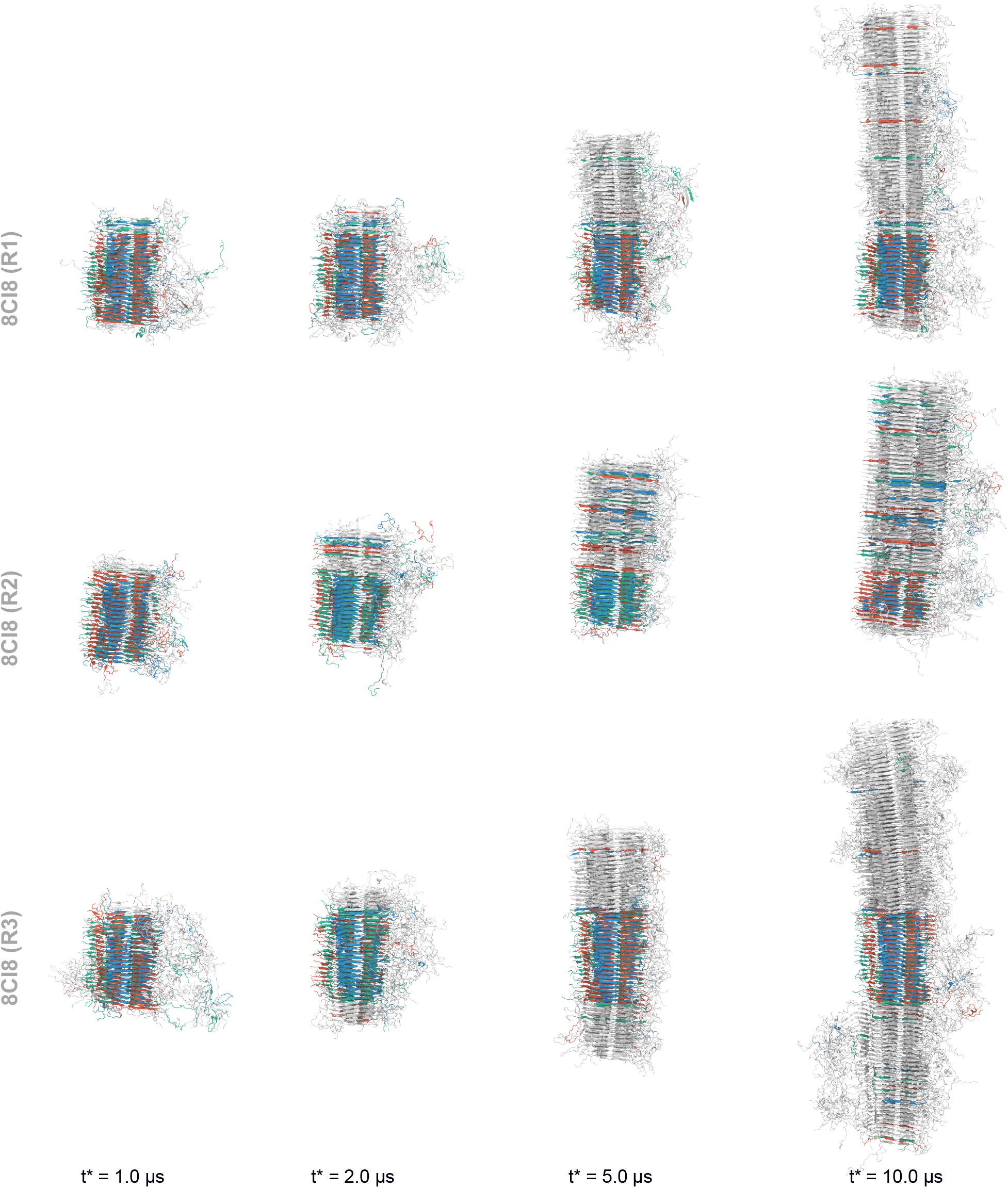
Representative snapshots of seeded fibril growth simulations of Nup98^FG^(298–327). Snapshots from three independent simulations, each initialized from a 30-layer fibril seed of Nup98^FG^(298–327) (PDB ID: 8CI8), are shown at 1.0 μs, 2.0 μs, 5.0 μs, and 10.0 μs. The three protofibrils of the original seed are color-coded in red, green, and blue, while newly attached monomers are shown in gray.

**Suppl. Fig. S10.**
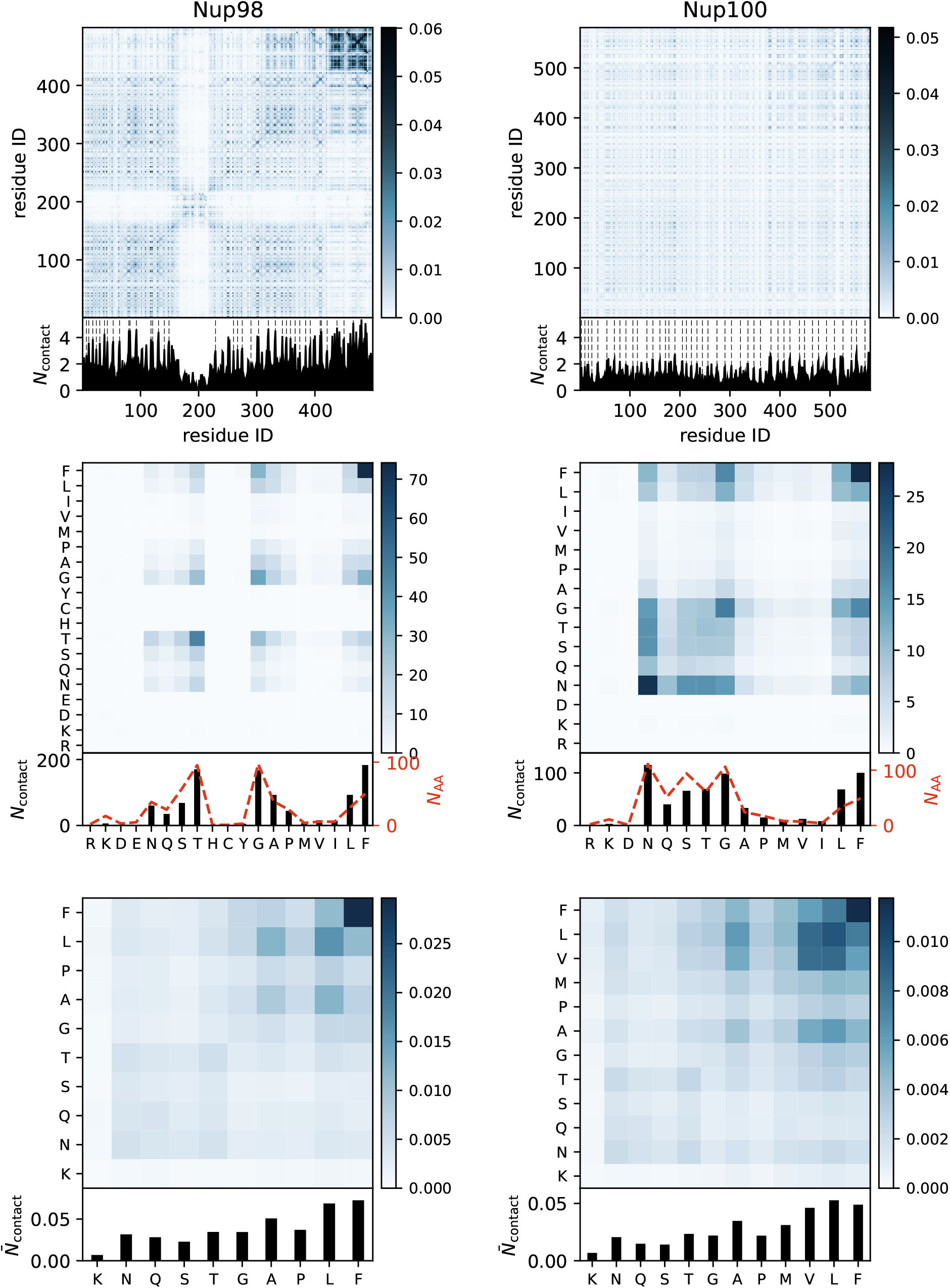
Intermolecular contact maps of the FG-Nup condensates of Nup98 and Nup100. (top) Average number of intermolecular contacts per protein replica as a function of residue number. The bottom figures show the one-dimensional summation, where the dashed lines indicate the location of the FG motifs. (middle) Average number of intermolecular contacts per protein replica as a function of residue type. (bottom) Average number of intermolecular contacts per protein replica as a function of residue type, normalized for residue occurrence in the protein sequence.

**Suppl. Fig. S11.**
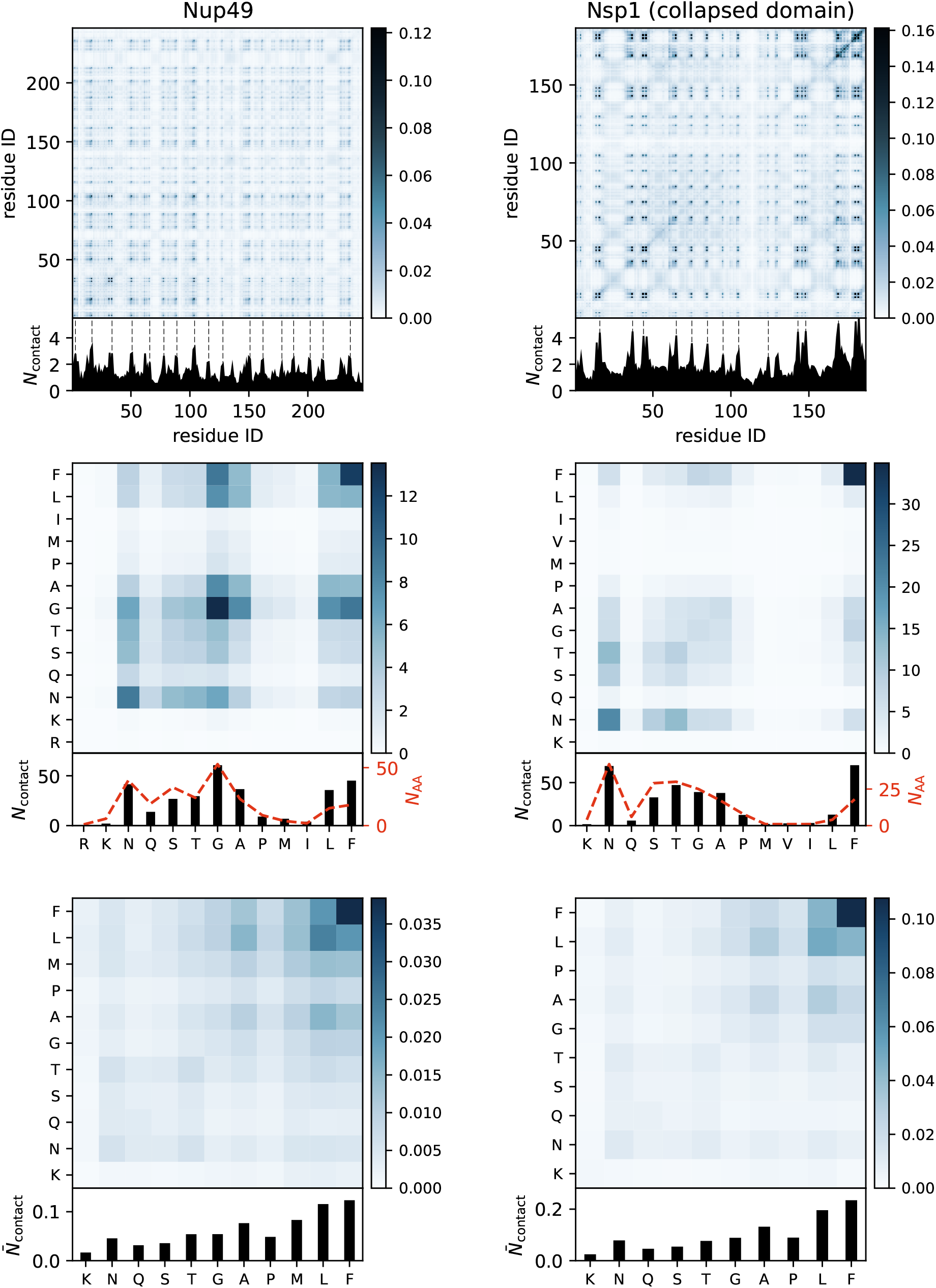
Intermolecular contact maps of the FG-Nup condensates of Nup49 and the collapsed domain of Nsp1. (top) Average number of intermolecular contacts per protein replica as a function of residue number. The bottom figures show the one-dimensional summation, where the dashed lines indicate the location of the FG motifs. (middle) Average number of intermolecular contacts per protein replica as a function of residue type. (bottom) Average number of intermolecular contacts per protein replica as a function of residue type, normalized for residue occurrence in the protein sequence.

**Suppl. Fig. S12.**
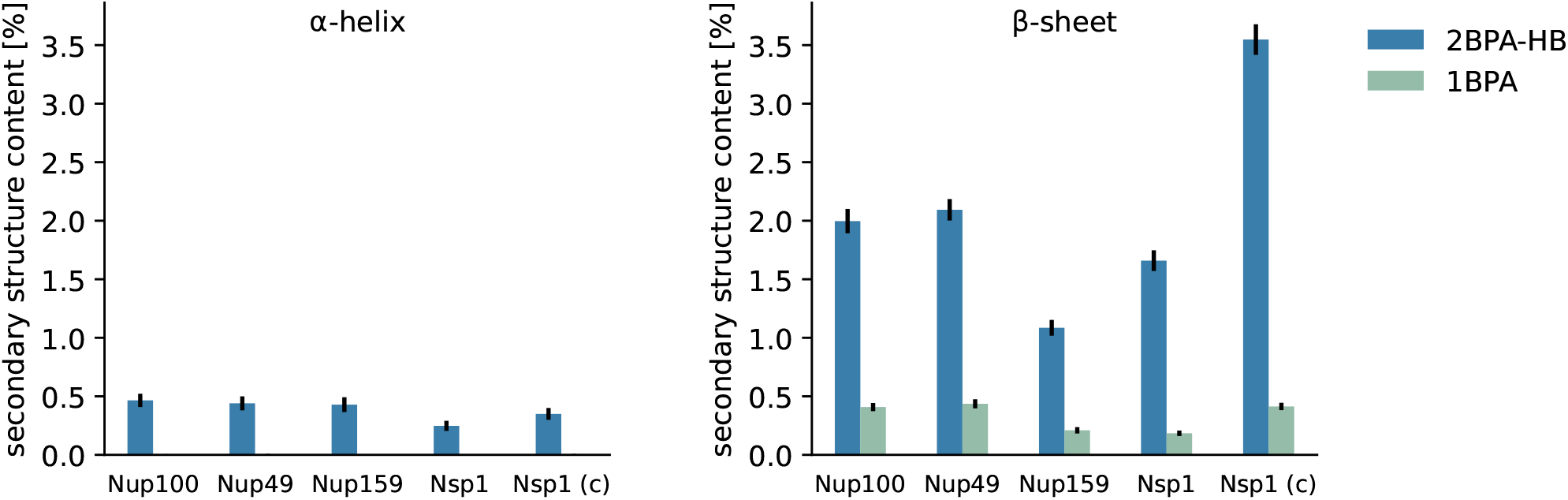
Comparison of secondary structure content between the 2BPA-HB and 1BPA models. Average α-helical (**left**) and β-sheet (**right**) content for phase separation simulations of five FG-Nup systems: Nup100, Nup49, Nup159, Nsp1, and the collapsed domain of Nsp1. Secondary structure content is defined as the fraction of residues adopting the respective conformation and was computed over the second half of each trajectory, with frames sampled every 50 ns. Results from the 2BPA-HB model are compared to corresponding simulations using the 1BPA model, where trajectories are taken from our previous work [10].

**Suppl. Fig. S13.**
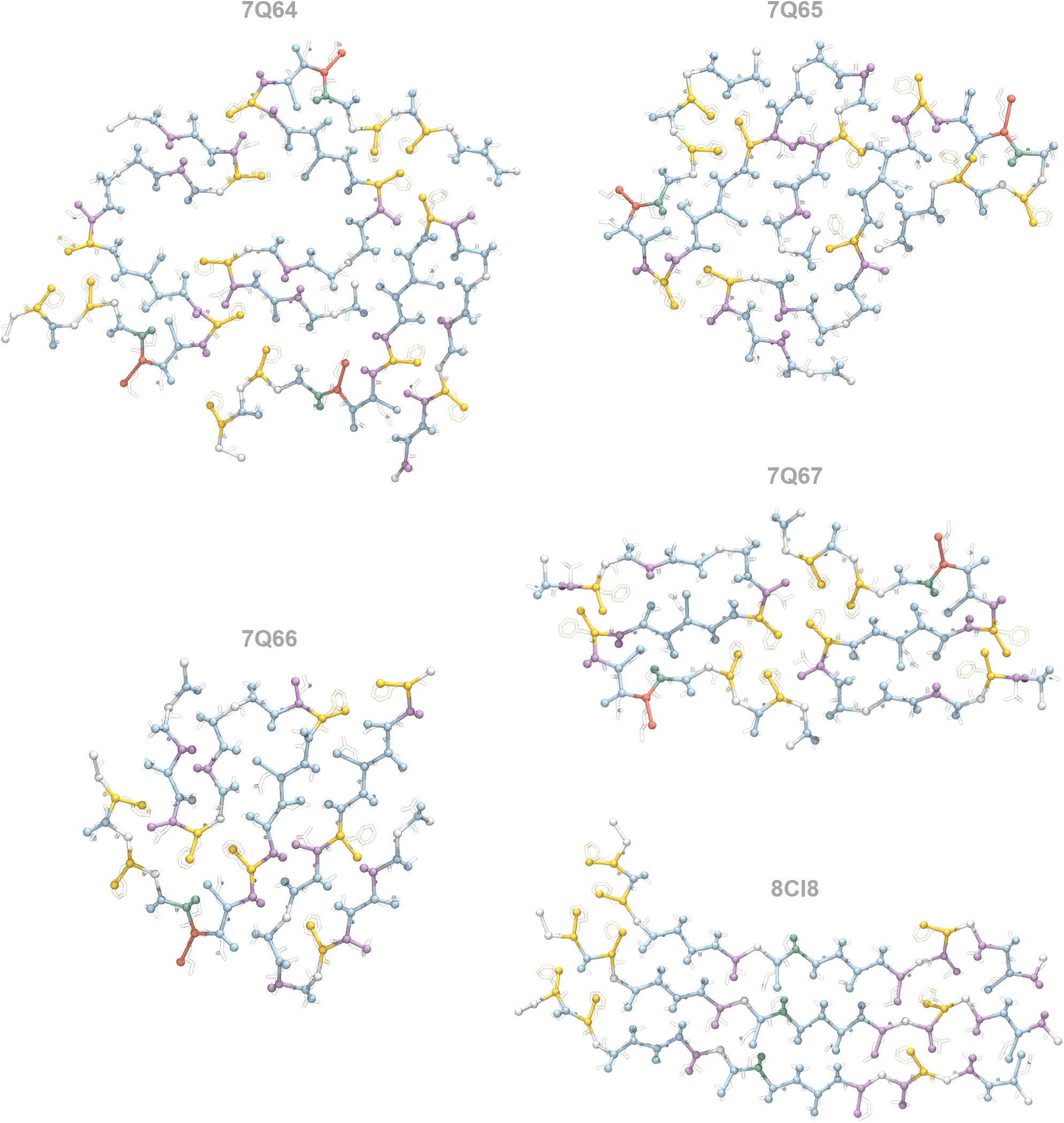
Comparison of coarse-grained and all-atom representations of Nup98 fibrils. For each of the five fibril structures used in this study, a single layer is shown in the 2BPA representation overlaid with the corresponding all-atom configuration (heavy atoms only, shown in transparency). Residues are colored by type: polar (blue), hydrophobic (purple), aromatic (yellow), cationic (red), glycine (white), and proline (green). Overall, the fixed side chain positions in the coarse-grained model align well with the corresponding all-atom interaction centers. Deviations observed for some residues reflect the conformational heterogeneity of side chains in the all-atom structures, which can adopt multiple orientations not captured by the static coarse-grained representation.

**Suppl. Fig. S14.**
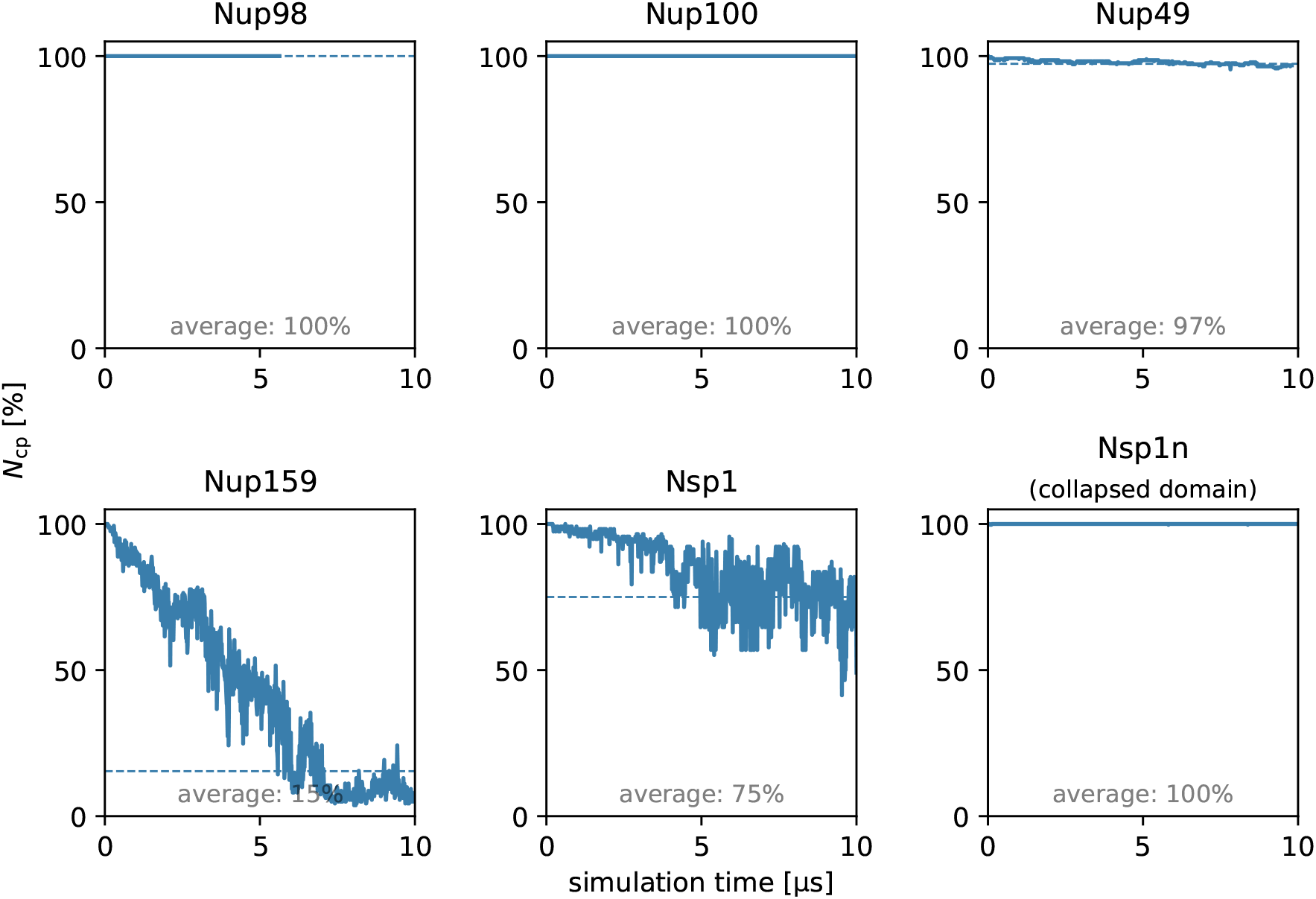
Convergence of phase separation simulations. Time evolution of the size of the largest molecular cluster, *N*_cp_ (%), for each phase separation simulation. Dashed lines indicate the average cluster size computed over the final 5 μs of each trajectory. All simulations were initialized from a preformed condensate, such that *N*_cp_ = 100% at *t*\* = 0. In all cases, the cluster size reaches a steady value within approximately 5 μs, indicating convergence to a dynamic equilibrium.

